# Highly porous granular hydrogels reinforced by electrospun hydrogel fibers for long-term stability

**DOI:** 10.1101/2024.10.28.620615

**Authors:** Jack Whitewolf, Lindsay Riley, Brooke Brady, Greg Grewal, Tatiana Segura, Chris Highley

**Affiliations:** Department of Biomedical Engineering, University of Virginia; Department of Biomedical Engineering, Duke University; Department of Chemical Engineering, University of Virginia

## Abstract

Granular hydrogels (GHs) formed from hydrogel microparticles (HMPs) possess inherent, microscale porosity. In GHs including microporous annealed particle (MAP) hydrogels as well as unannealed GHs, increases in pore size and total porosity correlate with enhanced cellular infiltration and growth. However, increased porosity or pore size are typically achieved by decreasing HMP density, reducing interparticle contacts and weakening hydrogel structure. Here, to achieve a highly porous GH without compromising stability, we developed an approach using sacrificial HMPs to achieve high, mesoscale porosity and high-aspect ratio hydrogel fibers to reinforce the scaffold. The use of sacrificial HMPs circumvented the need to decrease packing density, creating GHs with up to 60% porosity. The hydrogel fibers span particles in the porous space, stabilizing the porous structure by enhancing interactions among GH components. Upon 5-10% v/v fiber incorporation, highly porous GHs retained their structure over 28 days of *in vivo* culture, compared to 4 days in GHs without fibers. These scaffolds supported and promoted endothelial cell growth as porosity increased. Overall, this study presents a stable, highly porous granular scaffold system with high processability and cytocompatibility.

## Introduction

Hydrogels have been widely used as 3D cell culture platforms. Synthetic and natural polymers have been engineered to recapitulate characteristics of the extracellular matrix (ECM), including mechanical properties, degradability, and bioactive ligands that guide cell fate. A popular approach to hydrogel engineering is the fabrication of granular hydrogels (GHs), scaffolds formed from microscale building-block components^1^. Hydrogel microparticles (HMPs) can be packed to create GHs that transition from a solid-like state to a liquid-like state^2–5^. Specifically, when the applied stress is less than the yield stress of the granular system, the bulk material responds elastically to deformation; however, once applied stress surpasses the yield stress, the HMPs can flow past one another, and the bulk material exhibits shear-thinning behavior. This property is particularly desirable for injection-based applications and 3D printing^6–8^. In addition to their attractive flow property, packed HMPs form interconnected, micron-scale porosity among the particles, which primes the system for unhindered cellular infiltration^6,9,10^. This porosity is preserved even upon interparticle crosslinking, as shown prominently with microporous annealed particle (MAP) scaffolds^6,11^. HMPs that compose MAP scaffolds can be injected into wound sites and subsequently crosslinked to form a microporous scaffold, which has been shown to support tissue regeneration^6,11–14^.

Evidence suggests that further increasing the pore size and porosity in GHs enhances cellular infiltration^9,15^; however, altering these parameters can present many practical challenges. Currently, pore size and porosity (defined as non-particle volume fraction) in GHs are largely controlled by altering the size and packing density of HMPs. Increasing HMP sizes proportionally scales up pore size without affecting the porosity (Schematic 1A). However, increasing HMP size can impact packing behavior and affect the material’s bulk properties undesirably^5,9,16^. Additionally, the total scale-up in pore size eliminates smaller pores in the scaffold, which can result in cells settling at the bottom of the larger interparticle spaces, presenting challenges for *in vitro* culture^5,17^. To increase overall porosity of GHs, a common approach is to decrease HMP packing density^15^ (Schematic 1B). This can be accomplished by increasing interstitial fluid volume within the system. However, since the stability of GHs correlates with the number of contact points among HMPs, increases in interstitial volume that physical separate particles result in external forces more easily disrupting scaffold. Therefore, while increased porosity in GHs is desirable for cellular infiltration, it remains a challenging to do this without compromising the mechanical integrity of the scaffold^18^.

**Schematic 1.**
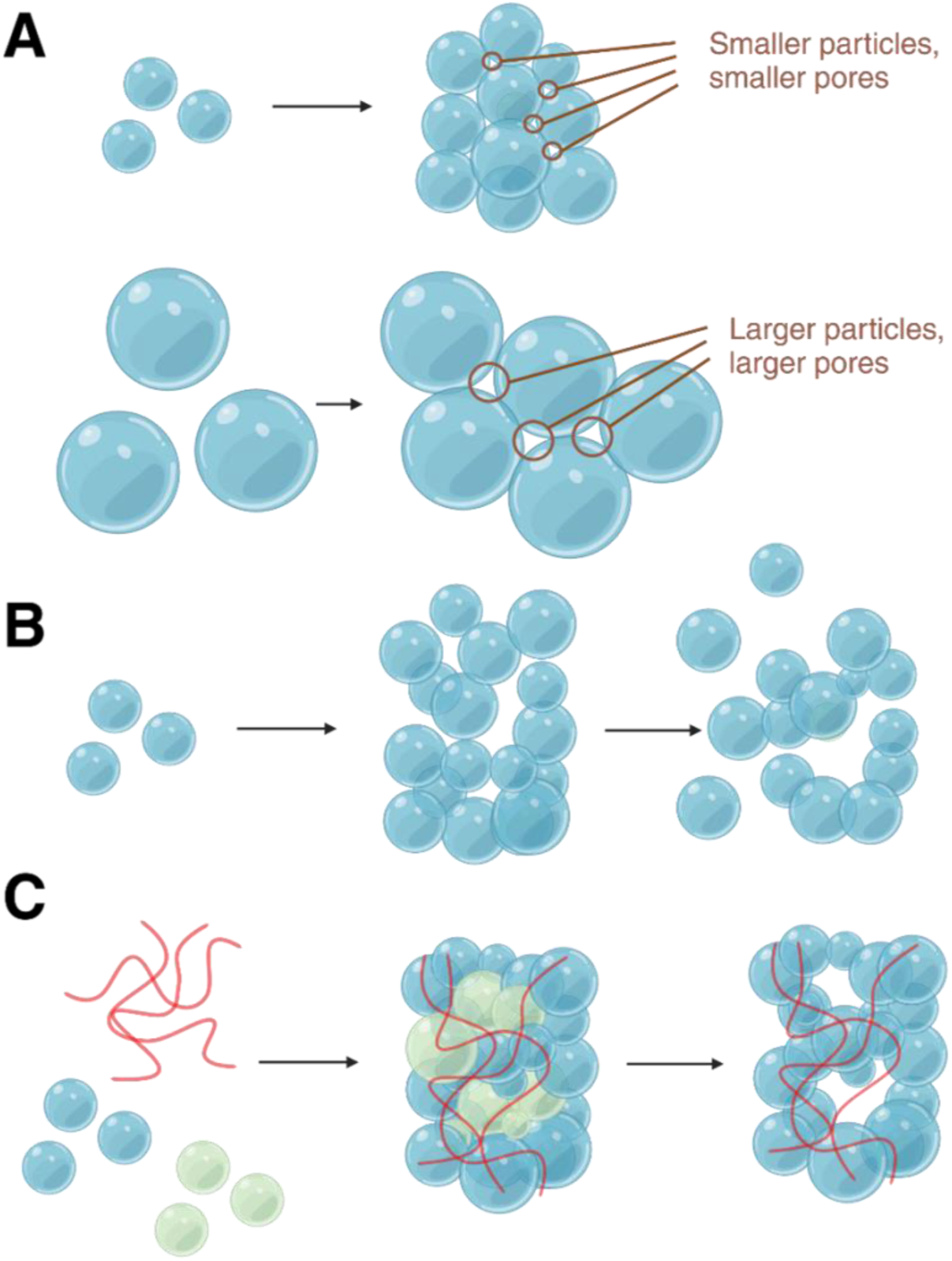
Approaches to increasing porosity in granular hydrogel scaffolds. **A)** Pores between particles scale with particle size: to increase pore sizes, large particles can be used. **B)** To increase porosity, particles can be less densely packed, but this reduces interparticle interactions and construct integrity. **C)** Pore sizes and porosity can be controlled precisely using removable particles, and porous constructs stabilized with hydrogel fibers.

Approaches to stabilize GHs that go beyond direct particle-particle crosslinking, as in MAP scaffolds, include crosslinked interstitial matrices or interpenetrating networks throughout the GH^18–22^. However, both methods require a polymer network to be formed within the interstitial space, which removes the microscale interstitial porosity critical to cellular infiltration. Alternatively, to improve GH stability, decreasing HMP size and increasing packing density both increase contact points, and decreasing particle stiffness can increase the total area of particle-particle interfaces. However, these options all produce smaller interstitial pore spaces that are less accessible to cells^15,23,24^.

In our work, we establish an approach to increase GH porosity while maintaining scaffold stability. Using sacrificial HMPs^25^, we demonstrate well-controlled porosity by changing the ratio of sacrificial HMPs to scaffold-forming HMPs. We show that interconnected mesoscale (dimensions 100 µm and greater) pores desirable for cellular infiltration can be achieved by design without using large HMPs or changing process parameters to achieve low particle densities. We introduce small volumes of high-aspect ratio hydrogel nanofibers into GHs to stabilize porosities that are not achievable in their absence (Schematic 1C). These long, thin fibers attach to HMP surfaces and thereby reinforce the particle network without significantly affecting total porosity. The fiber-HMP platform produces highly porous scaffolds with long-term stability that supports cell culture. We show that increasing porosity promotes cell growth in scaffolds, and that these materials can be injected with cells, laying a foundation for applications of fiber-reinforced GHs in cell delivery and regenerative medicine.

## Results and Discussion

### Fabrication and characterization of norHA HMPs

HMPs were generated from norbornene-modified hyaluronic acid (norHA). Hyaluronic acid (HA) is a naturally derived polysaccharide that is biocompatible, biodegradable, and amenable to chemical modification. The addition of norbornene pendant groups to HA enables reactions of norHA to thiolated molecules, as this highly efficient thiol-ene click chemistry allows for stoichiometric control over the consumption of norbornene groups by limiting the available thiol-containing groups (Figure 1A). This enabled the preservation of norbornene pendant groups for secondary conjugations. Here, we synthesized norHA with a 0.3 degree of substitution (DoS), confirmed by ^1^H NMR (Figure S1). By controlling the thiolated crosslinker to norbornene concentration, 0.3 DoS sufficiently supported both crosslinking required to form the HMPs and subsequent interparticle crosslinking within GHs to form MAP gels and fiber-HMP GHs (Table S1).

**Figure 1.**
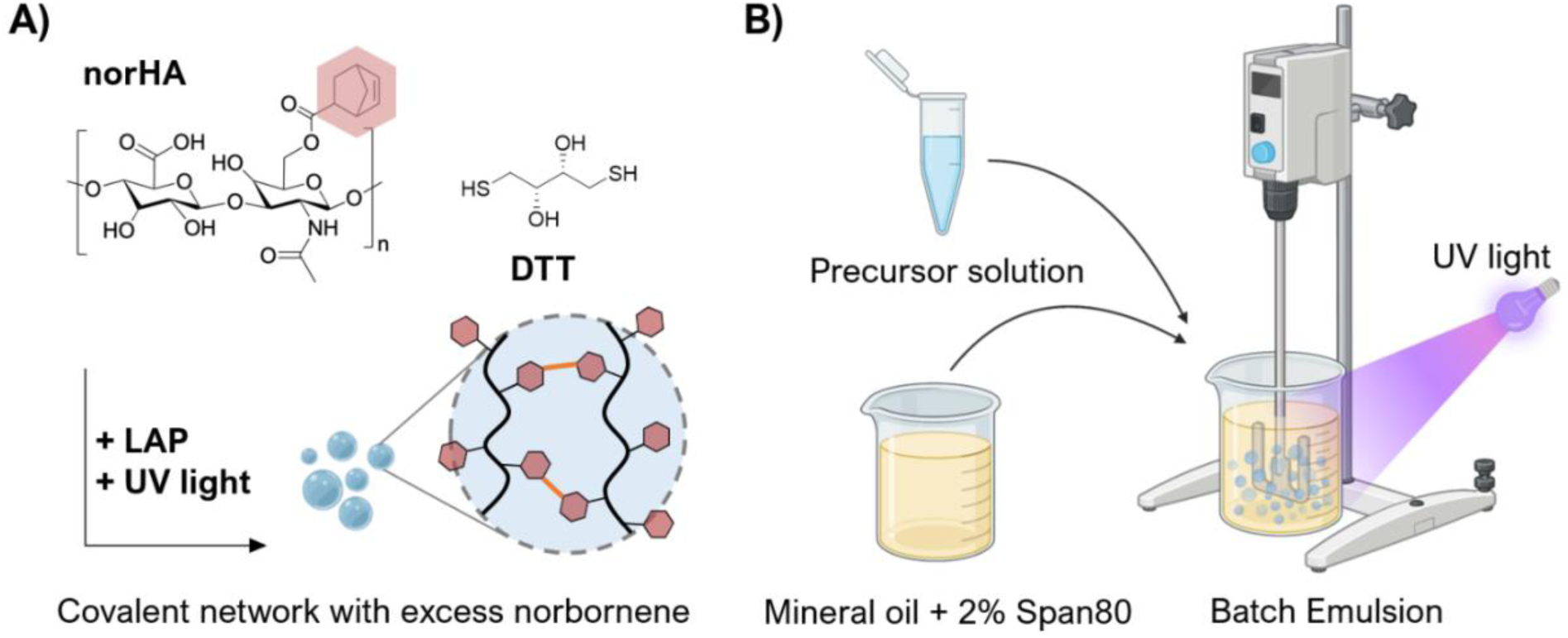
**A)** Thiol-ene click chemistry using norHA and DTT. Norbornenes conjugated to HA and DTT can be crosslinked under UV light with the presence of a photoinitiator (LAP). Stoichiometric control of DTT ensures there be excess norbornene for later reactions. **B)** HMP fabrication. HMP precursor (norHA, DTT, and LAP; aqueous phase) is mixed together with mineral oil with 2% Span80 (oil phase) and homogenized to form an emulsion. Application of UV light while stirring crosslinks the precursor solution into stable HMPs.

Using batch emulsification, we generated norHA HMPs (Figure 1B) with varying moduli and size by varying the HMP composition and the speed of homogenization, respectively. Soft, medium, and stiff HMPs were made using 2% w/v, 3% w/v, and 4% w/v norHA concentration and 2mM, 3mM, and 4mM DTT concentration, respectively (Table S1). The mechanical properties of the HMPs were characterized by measuring the storage moduli of bulk hydrogels with identical compositions. Soft, medium, and stiff HMPs had average storage moduli of 782 Pa, 1758 Pa, and 3797 Pa, respectively (Figure S2). We also altered the size of norHA HMPs by changing the spin speed of the homogenizer used for emulsification. Small, medium, and large HMPs were generated at 750 rpm, 1500 rpm, and 3000 rpm. For each modulus, we generated polydisperse populations of HMPs with mean diameters ranging from <10 µm to 50 µm (Table 1). HMP polydispersity increased as the stir speed decreased (Figure 2A); however, mean sizes were well-separated and statistically significant for groups across stir speeds. The effects of HMP composition on HMP mean diameter was negligible, due to the large coefficient of variation in our polydisperse populations (Table 1, Figure S2).

**Figure 2.**
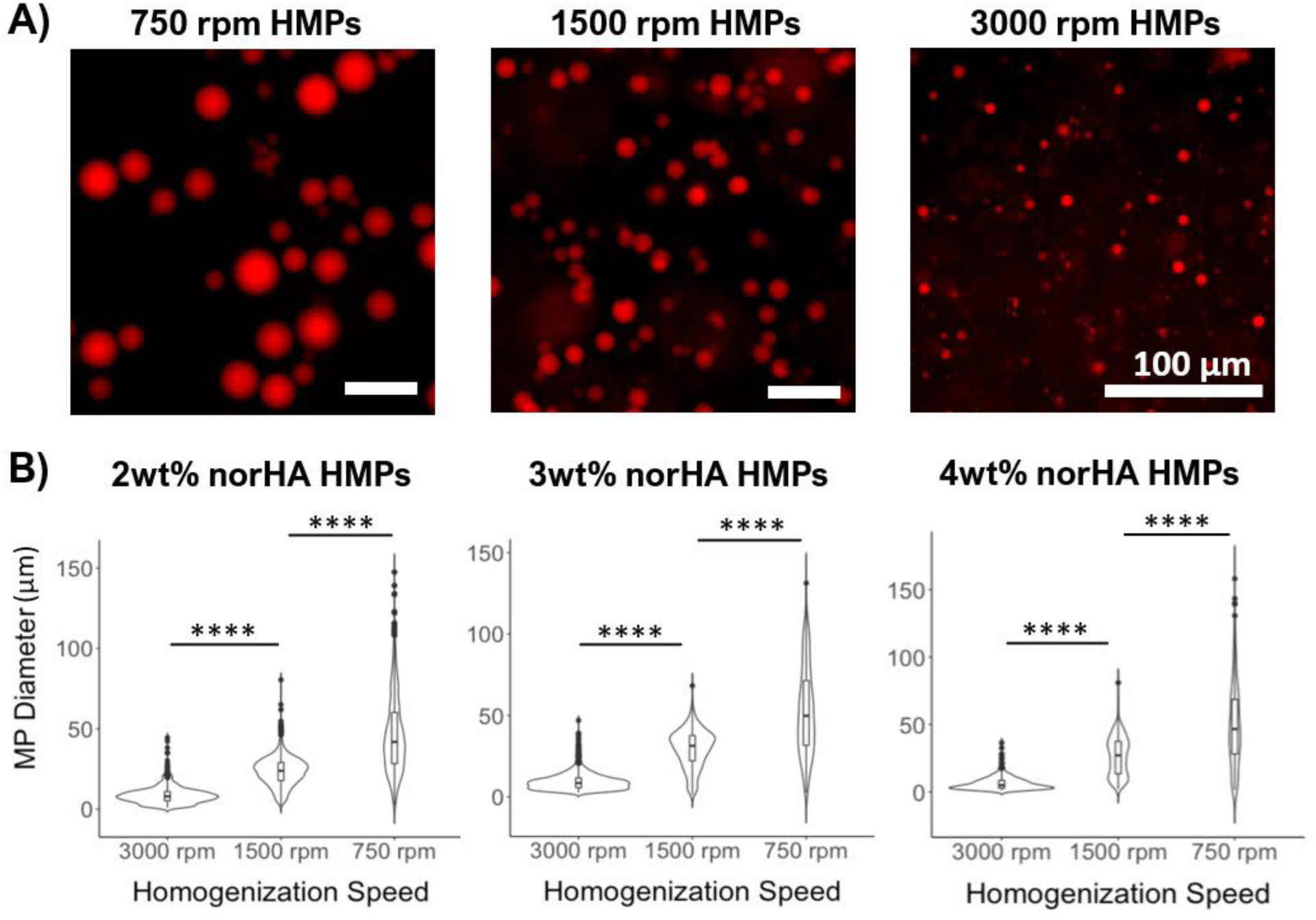
NorHA HMP size characterization. **A)** Size distributions comparing HMPs made from varying homogenization spin speeds. **B)** Fluorescent micrograph of HMPs at varying spin speeds. Scale bars = 100 µm.

**Table 1.**
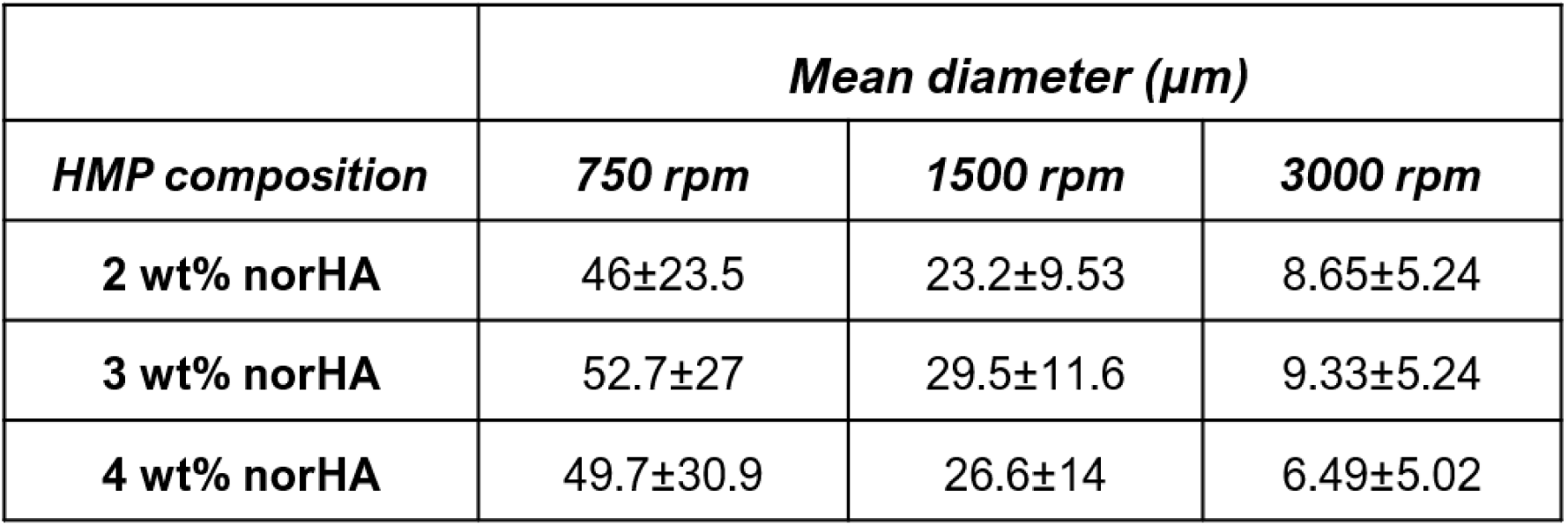
NorHA HMP mean sizes for all compositions and speeds.

Next, we conducted rheological tests to investigate the properties of GHs. GHs comprise HMPs that enter a jammed state when sufficiently packed^1–3^. Jammed HMPs exhibit self-healing behaviors where they can flow under external forces and recover when the applied force is removed. This property allows granular hydrogel formulations to be extruded from a needle, enabling various applications in injectable therapeutics and 3D printing. We confirmed that our jammed HMPs exhibit the characteristic self-healing behavior using time sweeps alternating between 0.5% and 500% strain (Figure S3). For all compositions and sizes, packed norHA HMPs exhibited solid-like behavior below the yield strain (0.5% strain) and liquid-like behavior above the yield strain (500% strain). To assess the bulk moduli of jammed HMPs at rest, frequency and strain sweeps were conducted for all HMPs. We observed that changes in granular hydrogel moduli correlated to changes in the moduli of the component HMPs. In considering the effect of HMP moduli on bulk moduli when size was held constant, we show that increases in HMP modulus resulted in increases in the modulus of the granular hydrogel as a bulk (Figure 3A, S3). However, changes in HMP sizes largely did not affect the bulk modulus, with one exception observed in the 4% norHA group (Figure 3A).

**Figure 3.**
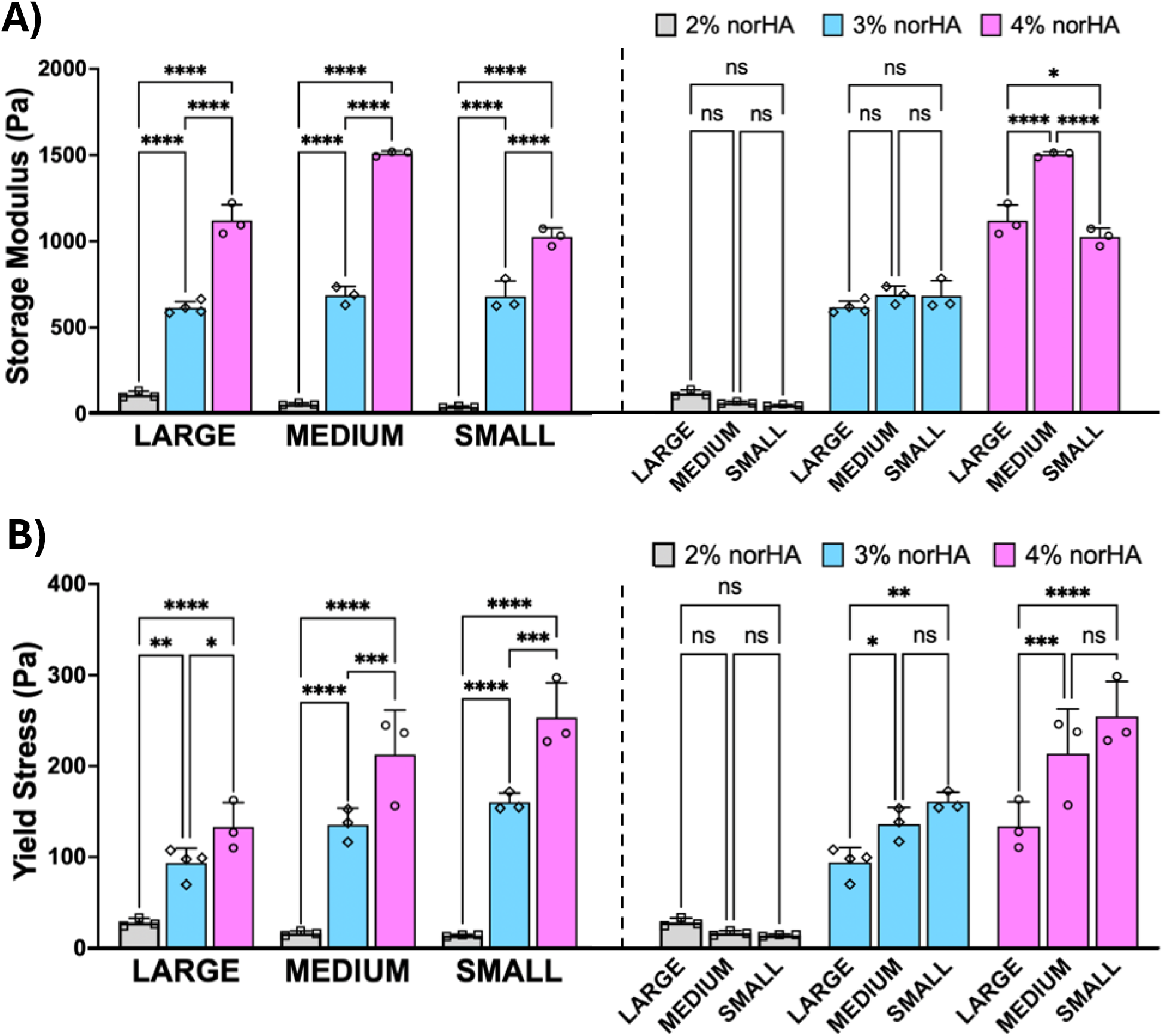
**A)** Quantified storage modulus of norHA HMPs for all compositions and sizes, compared based on composition (left) or size (right). **B)** Quantified yield of norHA HMPs for all compositions and sizes, compared based on composition (left) or size (right). All error bars denote standard deviations.

Another important feature of granular hydrogels is their resistance to flow, often described by the yield stress of the system^5,26–28^. Here, we measured yield stress using stress sweeps supplemented by shear rate sweeps focused on lower shear rates (Figure S4, Figure 3B). Yield stress of each formulation was quantified using results obtained from stress sweeps. Shear rate sweeps at low shear rates provided an indirect measure of shear stresses, which was used to confirm no significant deviations occur between measurements from either method. Similar to results obtained for bulk moduli, for HMPs of the same size, yield stresses of packed HMPs increased as the moduli for component HMPs increased (Figure 3B). We also observed that for 3% norHA and 4% norHA HMPs, yield stress increased as HMPs became smaller (Figure 3B). This finding largely corroborates literature findings of yield strain changes with respect to HMP sizes^6^.

Further, this trend was corroborated by qualitative experimental observations. 2wt% norHA HMPs, even when fully packed, flow very easily and could not hold their shape at rest, reflecting the low yield stress as well as storage modulus. On the other hand, small 3wt% norHA HMPs and all 4wt% HMPs presented challenges in processing: during pipette transfer, these HMP formulations frequently stuck to the side of the pipette tip, and the bulk granular material fragmented or separated into discrete pieces during aspiration or extrusion.

### Creating tunable porosity in HMP scaffolds using sacrificial gelatin HMPs

The use of sacrificial HMPs^25,29^ is an alternative method for creating porosity that does not rely on changing particle size or packing density. Here, we mixed known volumes of norHA HMPs (scaffold-forming) and gelatin HMPs (sacrificial) at defined ratios to control porosity in GHs. Gelatin HMPs were fabricated similarly to norHA HMPs through emulsification (Figure S5) and mixed with norHA HMPs to form solid scaffolds. The gelatin HMP component can be liquefied (or sacrificed) at standard culture temperature (37°C), making them easily removable to create pore spaces. The process is shown schematically in Figure 4A, where the annealing step forms a MAP hydrogel with sacrificial gelatin HMPs among covalently interlinked norHA HMPs. Removing gelatin HMPs by melting results in enhanced porosity with mesoscale features. To observe porosity characteristics created from this process, we first fabricated gelatin HMPs with varying sizes and generated scaffolds where gelatin HMP diameters are less than, equal to, or greater than norHA HMP diameters (Figure S6). For each size combination, we generated scaffolds with norHA HMPs and gelatin HMPs mixed at 100:0, 75:25, and 50:50 volume ratios. These scaffolds are then imaged with FITC-dextran as the interstitial fluid (Figure 4A) to assess porosity.

**Figure 4.**
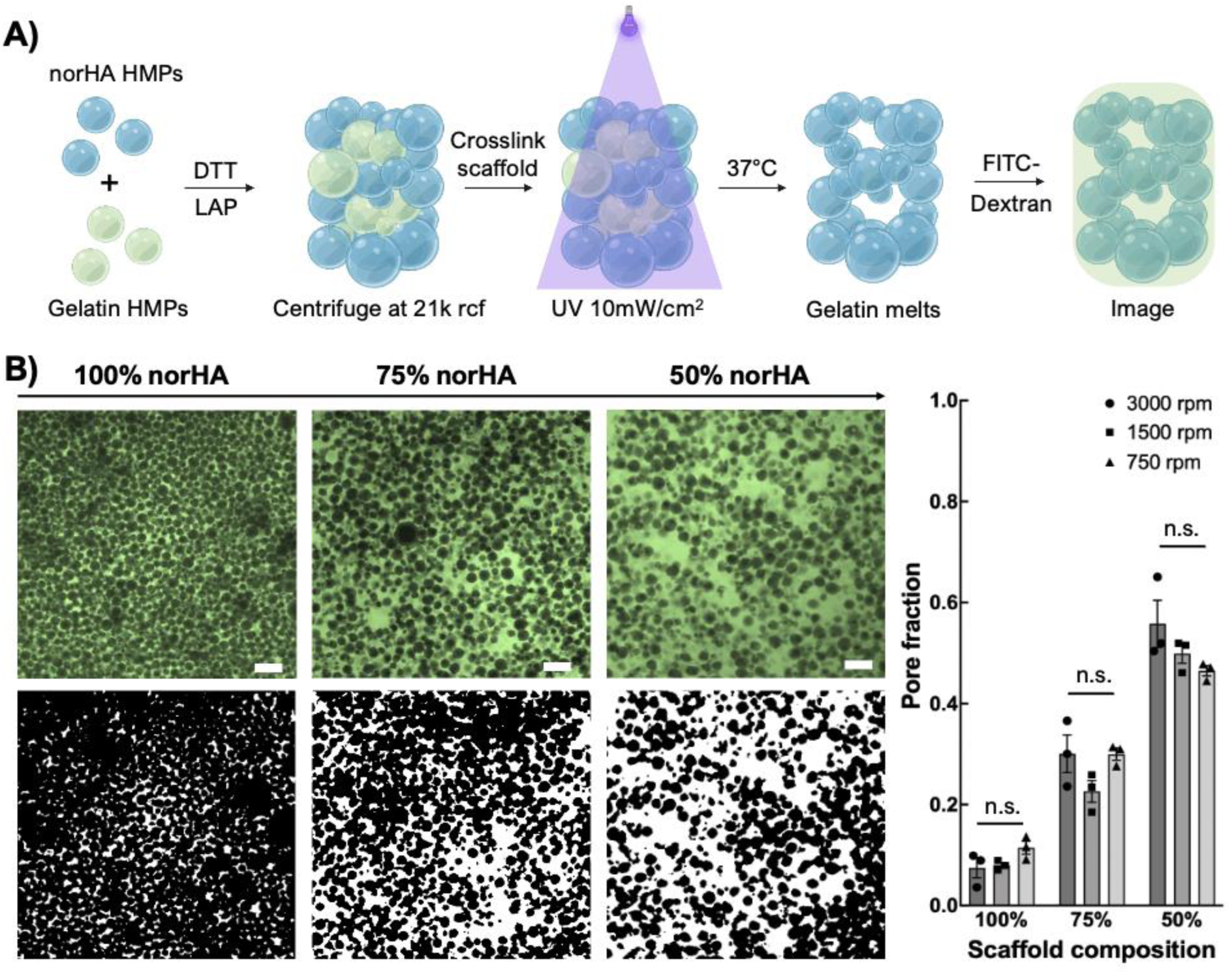
**A)** HMP scaffold fabrication. Desired ratios of norHA and gelatin HMPs, along with DTT and LAP, are combined and packed by centrifugation. The packed HMPs were then crosslinked and cultured at 37°C to liquefy gelatin HMPs. The scaffolds were then submerged in FITC-dextran solution and imaged. **B)** *Left:* Representative confocal images (top) and corresponding thresholded images (bottom) of scaffolds with varying ratios of norHA HMP (750 rpm) to gelatin HMP (2000 rpm). *Right:* Quantified void volume fraction for each scaffold composition. Scale bars = 100µm. Error bars denote standard

We observed that when gelatin HMP diameters were greater than norHA HMP diameters, more isolated pockets of void space formed where large sacrificial particles had previously resided. However, equal-sized or smaller gelatin HMPs increased void spaces more uniformly in scaffolds (Figure S6). The total porosity, however, was predominantly based on the ratio of gelatin-to-norHA HMPs and was only marginally affected by sacrificial HMP sizes. For subsequent studies, we created scaffolds using gelatin HMPs with diameters smaller than norHA HMP diameters to minimize the isolated pockets of void space in our scaffolds.

We then quantified changes in scaffold porosity based on incorporated sacrificial HMP ratios and norHA HMP sizes. For scaffold of each volume ratio (100:0, 75:25, and 50:50), we generated separate scaffolds using small, medium, and large populations of norHA HMPs to elucidate the effect of scaffold-forming HMP size on porosity. We found that for all norHA HMP sizes, we consistently generated scaffolds with ~10% porosity for 100% norHA scaffolds; ~30% porosity for 75% norHA scaffolds; and ~50% porosity for 50% norHA scaffolds, with negligible differences that are not statistically significant within groups (Figure 4B).

We then quantified pore sizes from 2D confocal slices using a custom MATLAB script that applies a standard connected-components approach to identify isolate pores and compute their area. For all scaffold compositions, we plotted pore area distribution by number frequency and confirmed that smaller pores make up the majority (Figure S7). As anticipated, we saw the presence of increasingly larger pores as the percentage of sacrificial HMPs increases (Figure S7). Figure 5A shows representative images of pore characteristics as porosity increased. For our scaffolds containing sacrificial HMPs, we observed that large, connected pores form the majority of the total pore area (represented in red) despite being fewer in number relative to small pores (represented in green). This is corroborated by plotting the pore area distribution normalized by total pore area for each scaffold group (Figure 5B). We saw that regardless of HMP sizes in the scaffold, the inclusion of sacrificial materials to obtain higher total porosity resulted in having larger pores that contributed to a majority of the porosity. This effect is most pronounced in scaffolds consisting of large norHA HMPs, where one or a few pores contributed to ~90% of the total porosity in the 50% scaffolds and ~60% of total porosity in the 75% scaffolds, indicating high degrees of interconnectivity.

**Figure 5.**
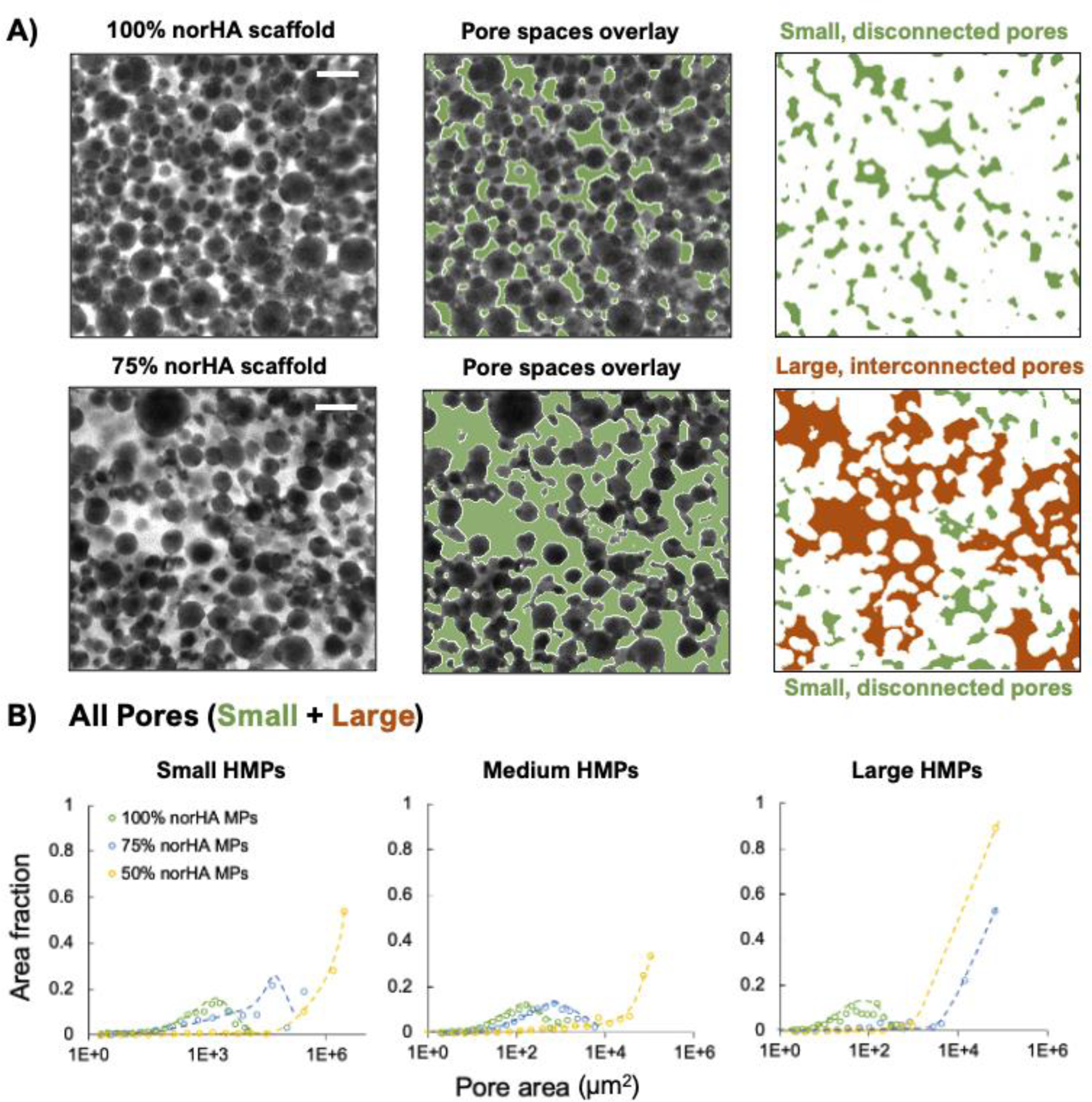
**A)** Visualization of 2D pore spaces. From left to right: confocal image of the scaffold with fluorescent interstitial (void) space; overlaying binary thresholding of void space (green); small pores (green) and large pores (red) separated from one another by white space. **B)** Pore area distribution normalized by total area. Small, medium, and large HMPs were used to make 100%, 75%, and 50% norHA scaffolds. For each distribution curve, pores from n = 3 corresponding scaffolds were sampled.

Further, we observed that in GHs formed with sacrificial HMPs, pore size distributions among the small pores were similar. To investigate this, we compared the distribution of only the smaller pores spaces in each group and found that the pore size distribution based on number fraction, as well as area fraction, was consistent among groups (Figure S8). This finding suggests that while higher porosity groups possess larger pores, smaller pores produced from random close packing of HMPs are still present.

The heterogeneity in pore sizes is unique to this system, where interconnected “mega” pores account for most of the porosity, yet many standard pores are present that are several magnitudes smaller. The co-existence of both types of pores allowed for the creation of non-uniform, multiscale void spaces that can be used to encourage cellular infiltration^30–36^.

### Computational assessment of 3D pore spaces in GHs

To extend 2D analysis of pore area distribution and general porosity to 3D considerations of pore volumes and interconnectivity, we utilized LOVAMAP^37,38^. LOVAMAP is a software designed for the analysis of pore spaces among packed particles. Using 3D images of the scaffolds, LOVAMAP can accurately discern 3D pore spaces and return information regarding pore volume and other pore characteristics. Here, we analyzed scaffolds composed of 3% w/v norHA HMPs fabricated at 750 rpm, as this formulation exhibited desirable flow properties and showed the highest connectivity across all porosity groups in our 2D analysis.

Using this formulation, we fabricated scaffolds with increasing porosity as previously described: 100% norHA, 75% norHA, and 50% norHA HMPs with the remainder composition being sacrificial gelatin HMPs. We fluorescently labeled norHA HMPs to allow for confocal imaging of our scaffolds. Reconstructed 3D images of the scaffolds were input into LOVAMAP to obtain visualizations and metrics of 3D pores (Figure 6A). Maintaining similar sampling volume for the scaffolds, we found that the total number of pores in the scaffolds decreased as scaffold porosity increased. At the same time, average pore volume increased as porosity increased (Figure 6B), indicating that larger but fewer pores are present as porosity becomes higher in our scaffolds.

**Figure 6.**
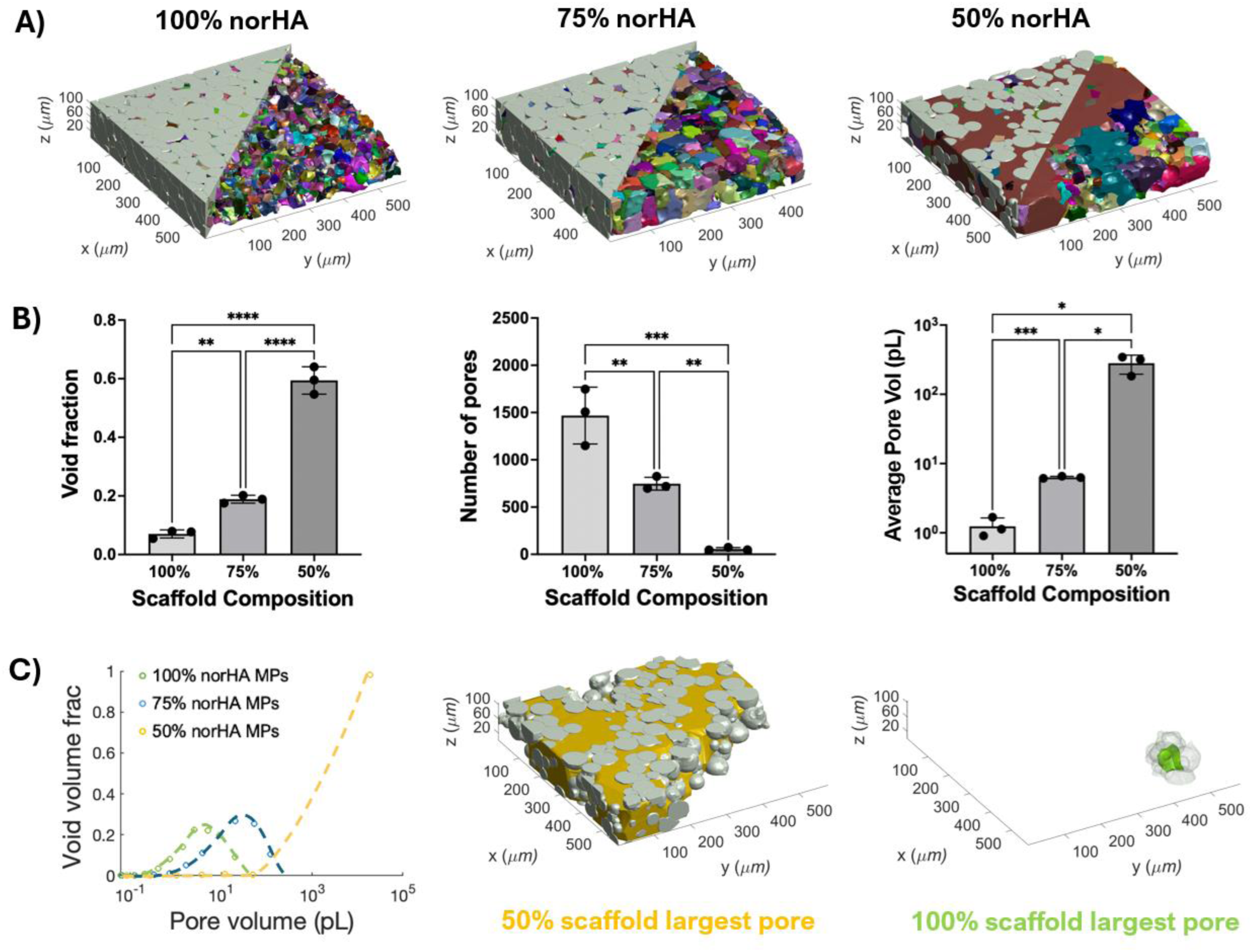
**A)** Representative LOVAMAP outputs of s with varying compositions. HMPs in grey; pores in color. **B)** from left to right: void fraction, total number of pores per scaffold, and average pore volume per scaffold. **C)** *Left:* pore volume normalized by total pore volume for each scaffold composition. *Middle and right:* representative largest pore for 50% norHA scaffold and 100% norHA scaffold. Error bars denote standard deviations.

Plotting pore volume normalized by total porosity showed the same trend as in 2D (Figure 6C), especially in our 50% scaffold where a single large pore accounted for over 90% of the total porosity. Plotting only small pores also showed similar distributions across groups (Figure S9), as seen in our 2D analysis. These results corroborate our previous findings about the porous heterogeneity of our system, where both ‘mega’ pores and small pores coexist.

Additionally, LOVAMAP analysis allowed us to differentiate between pores that interface with the surface of the sampled region (“exterior pores”) and those that are instead entirely enclosed within HMPs (“interior pores”) (Figure S9). While exterior pores represent open spaces that extend further into the scaffold, we can extrapolate to extension into the surrounding environment where tissue infiltration would occur. This is important toward understanding reciprocal access between the inside and outside of the scaffold through pores. LOVAMAP data showed that interior pore size was distributed similarly across groups, but the total number of interior pores decreased as scaffolds gained more porosity from sacrificial HMPs (Figure S9). For our least porous 100% norHA scaffolds, ~50% of the total pore volume came from interior pores, while in our highest porosity 50% norHA scaffolds, this number dropped to ~1%, indicating that most of the porous space was connected to the exterior (Figure S9).

Our pore analyses established that our system generates large, interconnected pores that offer easier access into and out of the scaffold, which may be desirable in tissue engineering (e.g., in supporting vasculature) and in regenerative medicine applications of MAP hydrogels and other GH systems. Importantly, porosity is a controllable feature that scales with the sacrificial HMP fraction. This circumvents the need to dilute packed HMPs or perform other processing steps to reduce packing density. Instead, mixtures can be ratiometrically combined to achieve predictable, high porosity in a single-step process. To further stabilize these HMP-based materials, we looked to quantify the impact of including hydrogel nanofibers in the interstitial space.

### Incorporation of electrospun fibers alleviates GH degradation

To generate hydrogel fibers for reinforcing HMP-based materials, we electrospun norHA fibers crosslinked by norbornene-DTT click chemistry. Crosslinked fibers were hydrated in PBS and segmented (Figure 7A) to yield high-aspect ratio fibers with a mean width of 1.6 ± 0.59 µm and a mean length of 95 ± 48.5 µm. Similar to HMP formation, in crosslinking electrospun norHA fibers, we controlled the norbornene to DTT ratio so that excess norbornene were present after crosslinking. The unreacted norbornenes on fibers enable secondary crosslinking with the norbornenes on HMP surfaces, strengthening fiber-HMP interactions.

**Figure 7.**
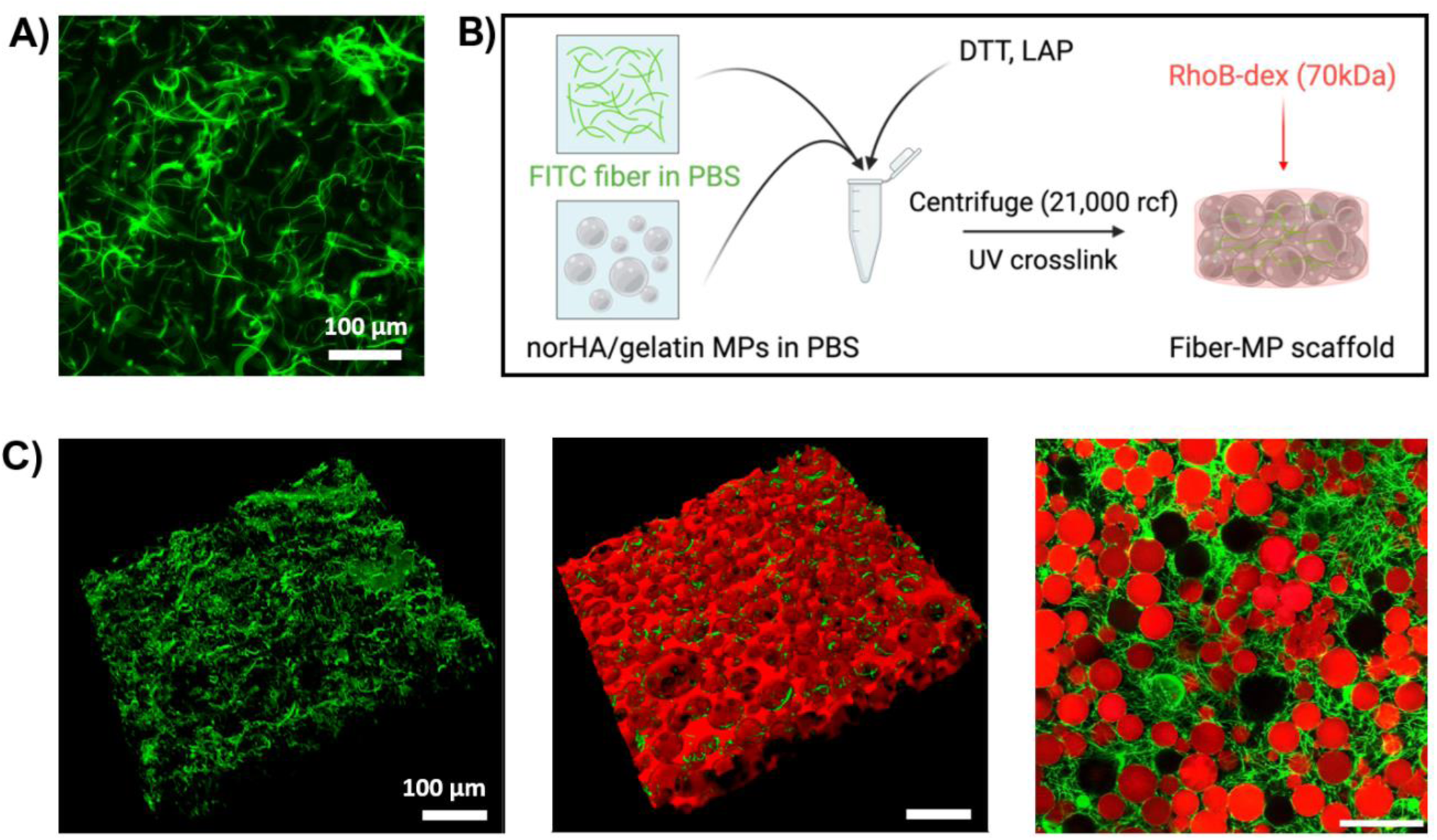
**A)** Representative fluorescent image of electrospun norHA fibers. A more dilute population of fibers was used for quantification of fiber sizes. **B)** Electrospun fiber incorporation into the scaffold. Fibers and desired ratios of norHA and gelatin HMPs were combined, along with DTT and LAP. The mixture is jammed through centrifugation and UV-crosslinked; then submerged in rhodamine B-dextran for imaging. **C)** *Left and middle:* 3D confocal image of fibers within the scaffold, and fibers combined with interstitial space (in red) within the scaffold. *Right:* Representative 2D confocal image of fibers (green) and norHA HMPs (red) in a 50% scaffold. All scale bars = 100µm.

We combined defined ratios of norHA fibers, norHA HMPs, and gelatin HMPs, to generate composite scaffolds with controlled compositions (Figure 7B). Confocal imaging of HMP scaffolds with fiber incorporation showed that fluorescently labeled fibers distributed within the interstitial space of the scaffolds (Figure 7C, left and middle), wrapping around volumes occupied by HMPs (spherical-shaped empty spaces), and we observed that fibers did not appear inside spaces occupied by HMPs. Additionally, fibers were well distributed throughout the space and could bridge between HMPs that were otherwise not in direct contact with one another (Figure 7C, right).

Next, we investigated the effects of fiber-based reinforcement on scaffold degradation, as a function of scaffold porosity. In our granular system, we had previously observed degradation by scaffold erosion with increasing porosity and concomitant drops in particle-particle contacts needed for annealing in MAP hydrogel systems and scaffold-stabilizing physical interactions in other GH hydrogels. Here, we added fibers to scaffolds of increasing porosity: 100% norHA HMPs, 75% norHA HMPs, and 50% norHA HMPs, and tracked their weight and appearance over 28 days. For each scaffold composition, we varied the fiber content (0% v/v, 1% v/v, 5% v/v, and 10% v/v) to elucidate the amount of fibers needed to reinforce granular hydrogels in long-term culture. As shown in Figure 8A, 100% norHA scaffolds retained ~70% of their initial mass without fiber reinforcement after 28 days of culture in PBS. Incorporation of 1%, 5%, and 10% fiber into 100% norHA scaffolds retained more mass (up to 80%) by the end of culture; however, there were no statistically significant differences, suggesting that these scaffolds are suitable for long term culture without fiber reinforcement. In 75% norHA scaffolds, the incorporation of fibers markedly enhanced the scaffold stability in long-term culture (Figure 8A). After the initial mass loss between day 0 and day 1 as the result of the removal of sacrificial particles, significant mass loss continued in scaffolds with 0% fiber. By day 7, only ~50% of the scaffold remained for 0% fiber scaffolds. Scaffolds with 1% fiber did not show significant differences compared to 0% fiber scaffolds for the first 6 days. However, from day 7 on, 1% fiber scaffolds degraded significantly less than 0% fiber scaffolds. On day 28, ~7% of 0% fiber scaffold and ~32% of 1% fiber scaffold remained. Neither group retained their shape at 28 days. In comparison, both 5% fiber and 10% fiber scaffolds held their shape over the 28 days of our study, and both retained >50% of their initial weight, which included the mass of the gelatin HMPs. Additionally, at no point of the study were the two groups statistically different from each other, indicating that both 5% fiber and 10% fiber scaffolds were suitable for maintaining stable GHs in long-term culture.

**Figure 8.**
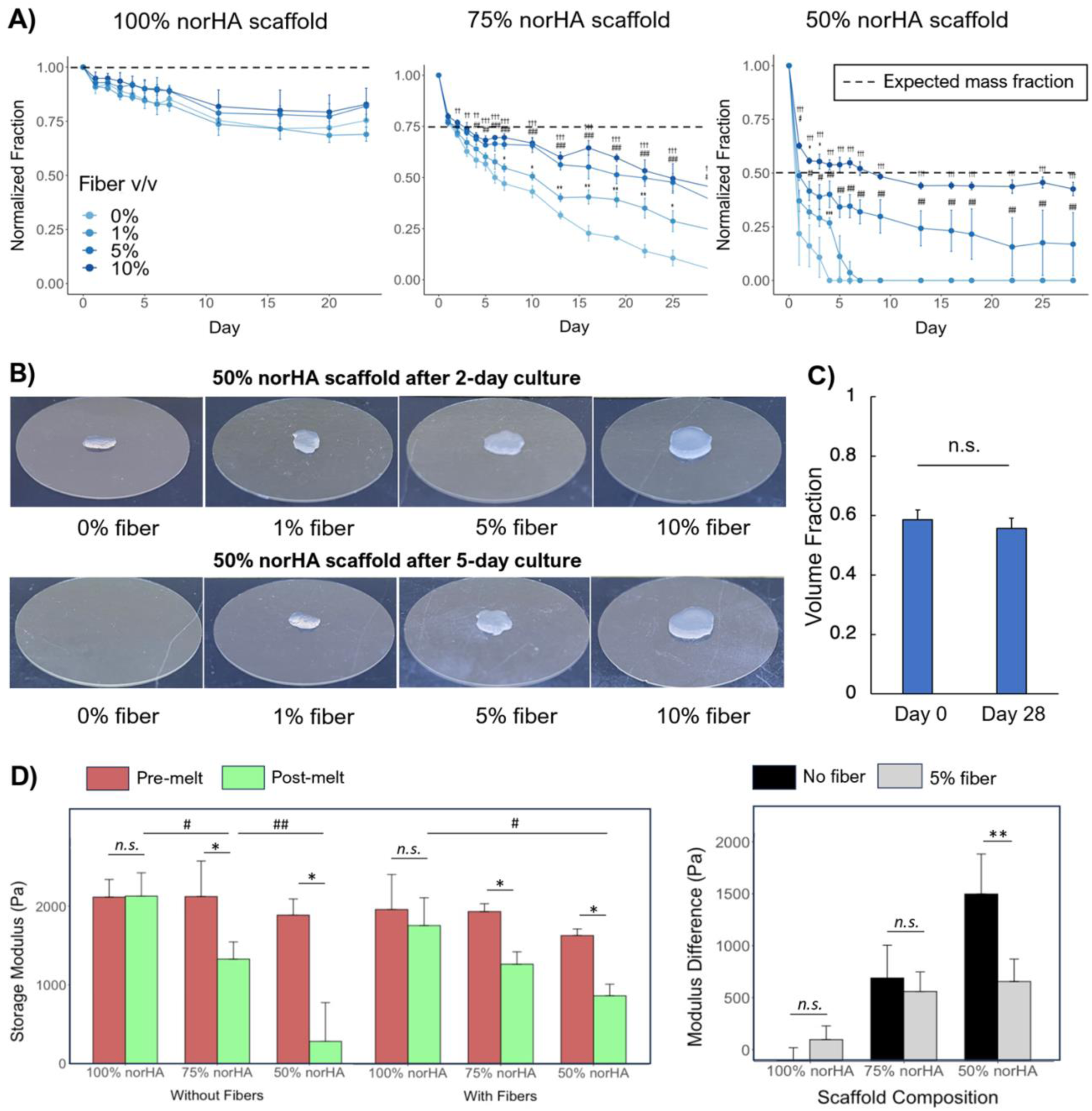
**A)** Remaining weight fraction of scaffolds tracked over 28 days normalized to scaffold weight at day 0. **B)** Visual presentation of effect of fiber amounts on 50% scaffold degradation over a period of 5 days. **C)** Porosity for 50% scaffold with 10% fiber at day 28 compared to day 0. D) *Left:* Rheological quantification of storage moduli of all scaffolds prior to and after gelatin HMPs liquefy. *Right:* Quantification of the moduli differences for each scaffold before and after gelatin liquefication. All Error bars denote standard deviations.

In our most porous scaffold group (50% norHA), the reinforcement effect of fibers was the most pronounced (Figure 8A, B). Without the incorporation of fibers, dissociation of 50% norHA scaffolds started by day 3, with scaffolds breaking apart into smaller pieces and losing most of their weight. Adding 1% fiber delayed the start of dissociation, but no scaffolds remained after 6 days. Incorporation of either 5% or 10% fibers in 50% norHA scaffolds was able to stop the scaffold from dissociating, with 10% fiber incorporation retaining significantly more scaffold mass after 28 days. Further, while both 5% fiber and 10% fiber incorporation were able to keep the scaffolds from dissociating, only 10% fiber scaffolds were able to maintain their shape over the course of the study, a representative image is shown in Figure 8B. Confocal microscopy of 50% norHA scaffold with 10% fiber on day 28 showed that the internal structure of the scaffold was preserved, with porosity comparable to newly made 50% norHA scaffolds (Figure 8C), demonstrating that the fibers both had minimal effects on porosity and the potential to preserve pore spaces within the highest porosity scaffolds over weeks in culture.

We also investigated whether fiber incorporation strengthened the mechanical properties of HMP scaffolds using non-destructive oscillatory rheology. We measured the shear modulus of 100% norHA scaffolds, 75% norHA scaffolds, and 50% norHA scaffolds, with no fibers or with 5% fiber to stabilize the scaffold.

Prior to gelatin removal, the storage moduli of all HMP scaffolds were statistically equivalent. The inclusion of fibers did not significantly alter the moduli of the scaffolds at this point. When gelatin HMPs were removed in 75% and 50% norHA scaffolds, we observed significant decreases in the moduli (Figure 8D). The inclusion of 5% fibers appeared to reduce the drop in moduli; however, this reduction was statistically significant only for 50% norHA scaffolds (Figure 8D). This difference, however, was partially due to some 50% scaffolds falling apart during testing, and thus had recorded moduli of 0. Combined, these data suggest that low amounts of fiber incorporation do not alter scaffold bulk modulus, and that fibers work to passively tether the HMPs together rather than creating structures with increased mechanical robustness.

### Porosity in fiber-HMP scaffolds support and facilitate HUVEC growth

To assess potential for our fiber-HMP scaffolds to support cell growth in regenerative medicine and tissue engineering applications, we introduced HUVECs into these systems, looking towards the potential for creating vascularized 3D hydrogel systems. HUVECs were cultured within 100% norHA, 75% norHA, and 50% norHA RGD-modified scaffolds for 4 days (Figure 9A). Upon cytoskeletal and nuclear staining, we saw that in all scaffold groups, HUVECs were able to adopt elongated morphologies (Figure 9B). Additionally, increasing porosity generally increased the total cell number per scaffold volume, with a significant increase in cell number between 100% and 75% norHA scaffolds (Figure 9C).

**Figure 9.**
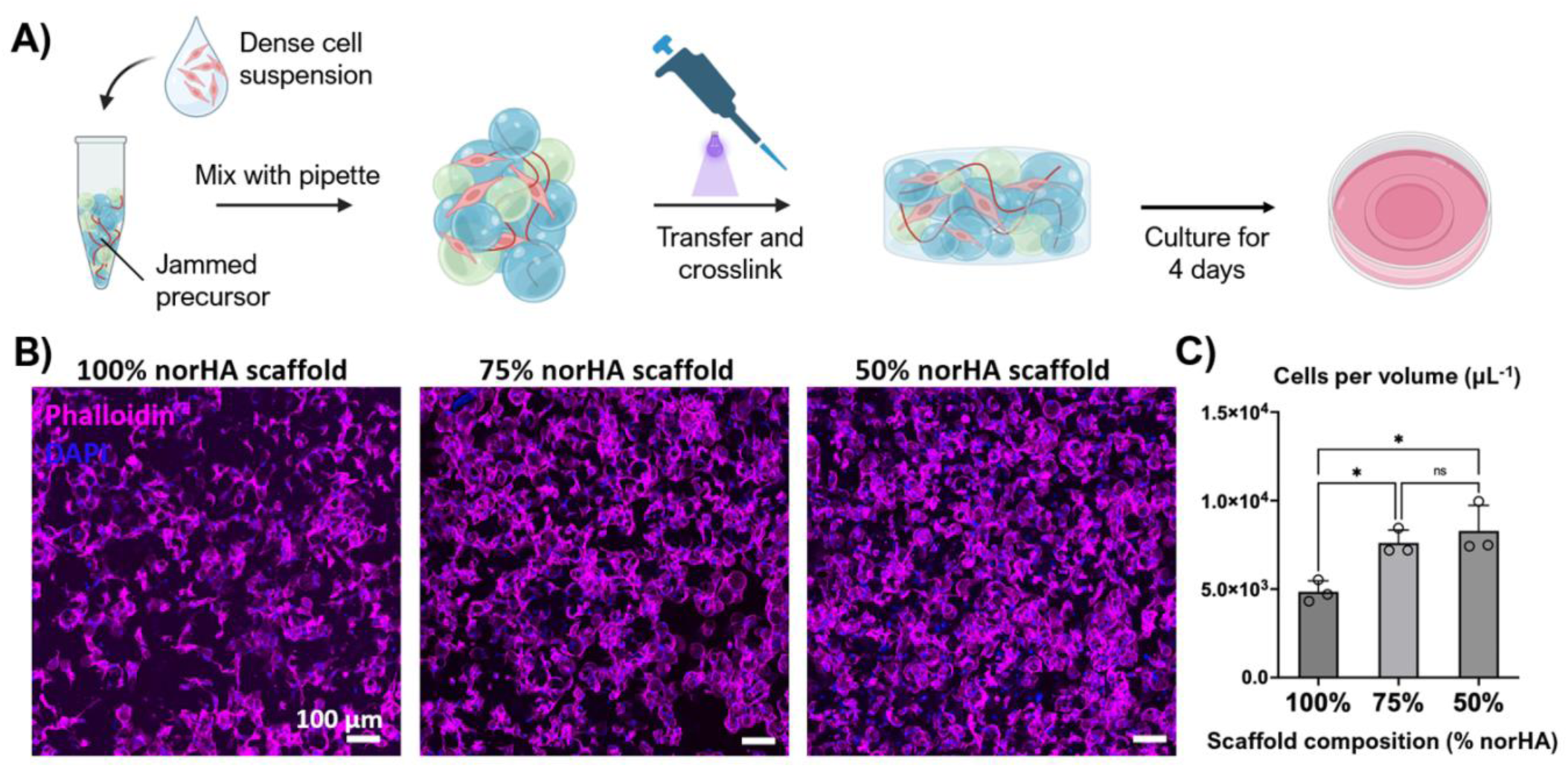
**A)** The process for HUVEC incorporation and scaffold formation. **B)** Representative max projection images of HUVECs cultured in fiber-HMP scaffolds with varying porosity **C)** Quantified cell number normalized by scaffold volume in each scaffold composition. Error bars denote standard deviations.

Additionally, we explored the injectability potential of our scaffold system. We extruded packed, 100% norHA HMPs with HUVECs included among the HMPs (Figure S10A), and subsequently crosslinked the HMPs into a scaffold. Viability assessments were performed (1) prior to the extrusion process, (2) immediately after extrusion and UV cross-linking, and (3) after three days of culture, to assess the effect of individual processing steps on cell survival. We observed high degrees of viability (80%-90%) throughout the process, with a slight reduction in cell viability after culture (Figure S10B, C). This might be attributed to delayed effects of the extrusion process as well as the lower porosity in 100% norHA scaffolds.

Together, our results indicated high cell viability and robust cell growth that increased as porosity became higher in fiber-HMP scaffolds. The ability of fiber-HMP scaffolds to support endothelial cells in culture suggests the potential for creating robust 3D cell cultures as well as systems that might ultimately contain engineered vasculature. Moreover, our granular hydrogel formulation can support cell viability through extrusion and might serve as a platform for cell delivery. We believe that our approach to controlling porosity will extend what is possible in engineering granular systems for use in 3D bioprinting and regenerative engineering applications.

## Conclusion

Creating high porosity in GHs has been challenging due to the associated reduction in particle packing and scaffold stability. Here, we have demonstrated a new approach for achieving high porosity in GHs while maintaining injectability and long-term stability. Through the application of sacrificial gelatin particles and scaffold-supporting hydrogel fibers, we demonstrated the fabrication of HMP-based hydrogels with porosity up to 60%, with *in vitro* culture stability for at least 28 days. We showed that our high-porosity scaffolds contain large, interconnected ‘mega’ pores formed from sacrificial particles but preserve local regions of close particle packing established during injection. When we formulated fiber-HMP hydrogels that incorporated 5-10% v/v electrospun hydrogel fibers, we were able to culture cells on scaffolds that were up to 60% porous. Without fiber reinforcement, these scaffolds would have dissociated in <4 days. Using fiber-reinforcement scaffolds, HUVECs showed improved proliferation as porosity increased. Combined, the methods and results in this paper present a particle-based hydrogel system with promising potential for *in vivo* regeneration and tissue engineering. Beyond our specific system, fiber-reinforcement offers a solution to the challenges of increasing the porosity in GHs, as well as increasing the stability by enhancing interactions among particles that form GHs and MAP hydrogels. This strategy, supplemented with the versatile modification potentials of individual HMPs or entire GH systems, may enable further advances in the rapidly expanding applications of MAP hydrogels and GH materials.

## Experimental methods

### NorHA synthesis and characterization

norHA was synthesized through a previously described method^39^. Briefly, HATBA was made by first dissolving sodium hyaluronate (lifecore biomedical) in water and adding Dowex® 50WX8 hydrogen form (Sigma). After filtering the resin at the end of the ion exchange, tetrabutylammonium hydroxide (Fisher) was used to titrate the filtered solution until pH reaches 7. Finally, the HATBA was frozen and lyophilized. To synthesize norHA, HATBA and 4-Dimethylaminopyridine (Sigma) were dissolved in DMSO under anhydrous conditions. 5-norbornene-2-carboxylic acid (Sigma) was then added to the solution. After the reagents became well mixed, di-tert-butyl dicarbonate (Sigma) was added and the temperature raised to 40°C. The reaction was allowed to proceed overnight and terminated by quenching with cold water. The mixture was purified through dialysis and precipitation, then lyophilized. ^1^H NMR was done to characterize the synthesized norHA by dissolving lyophilized norHA at 10mg/mL in D_2_O. NMR was conducted using a 600MHz NMR machine (Varian, Inova).

### NorHA HMP fabrication

norHA was dissolved at the desired weight percentages in PBS (2 wt%, 3 wt%, and 4 wt%). Then, 60mM LAP (Sigma) in PBS was added to adjust the final LAP concentration to 6mM. DTT (Sigma) was also added to reach final concentrations of 1mM of DTT per 1 wt% of HA in solution. After fully mixing, the solution was added to light mineral oil (Fisher) with 2% Span 80 (Sigma) at a ratio of 1:9 (solution : oil). The oil-solution mixture was then emulsified for 3 minutes at desired speed, then UV crosslinked for 15 minutes under stirring.

To retrieve the HMPs from oil and eliminate surfactant, the HMP-oil emulsion were first centrifuged at 3,000 rcf for 2 minutes. After discarding the oil, the HMPs were resuspended in hexane and centrifuged at 3,000 rcf again for 2 minutes. This was repeated twice. After the hexane wash, the HMPs were then resuspended in isopropanol and centrifuged at 1200 rcf for 5 minutes. The isopropanol wash was repeated twice to wash off the remaining hexane and sterilize the HMPs at the same time. After discarding the excess IPA, the norHA HMPs were resuspended and rehydrated in PBS. This process can be done sterilely to generate culture-ready particles. Finally, the HMPs were filtered through a cell strainer (Pluristrainer, Fisher) with mesh sizes ~1.5x the largest diameter HMP in the distribution to eliminate particle clusters formed through aggregation or faulty crosslinking.

### Gelatin HMP fabrication

Gelatin from bovine skin, type B (Sigma) was dissolved in PBS at 10wt% at ~40°C. The solution was then added to light mineral oil with 2% Span 80 (also heated to ~40°C) and emulsified at desired speed for 3 minutes. The emulsion was then allowed to cool to room temperature to form HMPs. Once cooled, the gelatin HMPs were washed and processed in the same way as norHA HMPs.

### Size characterization of norHA HMPs

norHA HMPs suspended in PBS at 1:1 volume ratio were centrifuged at 21,000 rcf for 5 minutes. After discarding the excess PBS, 2mM of thiolated peptide-rhodamine B (sequence GCGKKKG-RhoB) suspended in PBS was added at equal volume to the HMPs to achieve a final concentration of 1mM rhodamine B. 10mM LAP was also added to the solution to reach a final concentration of 1mM LAP. The HMPs were then vortexed with the solution so that particles are sufficiently suspended. The suspension was then placed under UV light for 5 minutes at 10mW/cm^2^. After conjugation, the HMP suspension was centrifuged at 21,000 rcf for 5 minutes again. The excess solution discarded and the HMPs were resuspended in fresh PBS. This washing step is repeated until no color was seen in the PBS after centrifugation.

The now fluorescent HMPs were imaged using a Leica DMi8 widefield microscope. HMPs were first diluted 1:100 volume ratio in PBS. Then 200μL of the suspension was added to a glass-bottom confocal dish and imaged on the microscope. Further dilution was carried out as needed so that HMPs do not overlap each other in the final micrograph. The image was then processed in FIJI, which delineated and analyzed the size of each particle. Plotting and statistical analysis was done using Rstudio. Welch’s ANOVA and Games and Howell post-hoc test was done for all hypothesis testing.

### Rheological tests for HMP flow properties

A DHR-3 rheometer (TA instruments) was used for all rheology testing. For all flow property assessments, the HMP samples were conditioned by pre-shearing at 100 s^−1^ for 10 seconds. Samples were then allowed to equilibrate for another 2 minutes before conducting measurements. All gap sizes are set to be 10 times greater than the larges HMP size in the population. Rheology tests are conducted on a Peltier plate, with a 20mm sandblasted solvent trap parallel plate geometry and temperature control at 20°C. For frequency sweeps, after conditioning, 1% strain was applied at frequency ranging from 0.01 Hz to 10 Hz in logarithmic increments. For strain sweeps, frequency was kept at 1 Hz, with strain varying from 0.01% to 500%, also in logarithmic increments.

### Fabricating porous norHA scaffolds

To create norHA scaffolds with varying porosity, both norHA HMPs and gelatin HMPs were first suspended in PBS at 1:1 volume ratio. From here, desired amount and ratio of both populations were calculated, and the appropriate amounts were mixed together. 10mM LAP and 25mg/mL DTT were added to the mixture to achieve a final concentration of 1mM for both. This mixture was then centrifuged at 21,000 rcf for 5 minutes; excess solution was discarded.

To crosslink the HMPs, desired amount HMPs from the last step was transferred with a wide-bore pipette tip. The HMPs were then UV crosslinked for 1 minute at 10mW/cm^2^. For HMP scaffolds with gelatin HMPs incorporated, PBS were added to cover the scaffold, and the scaffolds were cultured in a 37°C, humidified incubator for 20 minutes to liquefy the gelatin HMPs.

### Porosity characterization of norHA scaffolds

All norHA scaffolds were fabricated as described above. After scaffold formation, all scaffolds are submerged in 1mg/mL FITC-dextran in PBS and cultured at 37°C for >20 minutes. These scaffolds were then imaged under a Leica confocal microscope (Stellaris 5).

To assess the porosity of each scaffold, at least one and up to four randomly sampled regions were chosen for each scaffold. For each region, z-stack images were taken to penetrate 100μm into the scaffold, with step size no larger than 5μm. The images were thresholded with FIJI and the pore fraction for each slice calculated. The final porosity was estimated by the average of each z-stack image. If multiple regions were selected, the porosity from all z-stack images were averaged to be the final porosity of the scaffold. Replicates of at least 3 were done by fabricating a new scaffold for each replicate and repeating the above process.

### Pore space 2D analysis

To analyze the pore spaces in porous scaffolds, the same microscopy images were used as those for characterizing porosity. After thresholding in FIJI, the pixel values (0 or 256) and their corresponding XY coordinates were exported as a .csv file. To identify 2D pores, an in-house MATLAB script was developed to find connected pixels using pixel value and position data. The area of each pore space was then calculated in MATLAB. To classify small versus large pores, we used the largest pore found in 100% norHA scaffolds as the threshold cut-off for small pores.

Area fraction was found by first defining number of bins desired for graphic representation. Then, area fraction was calculated by dividing the sum of the pore areas in the same bin by the total pore area. The average pore area of each bin was used as the x-axis value in the distribution graph. To generate accurate representations of pore area distributions, we down sampled our z-stack images. We used at least three z-stack images, located at least 20 microns apart (typically z = 0 µm, z = 50 µm, and z = 100 µm) to capture and represent porosity located deeper within the scaffolds. For each distribution, at least three replicates of the corresponding scaffold were sampled in this fashion.

### LOVAMAP analysis of 3D pore space

norHA HMPs used for imaging was made fluorescent using AF 430 tetrazine (Lumiporbe). Briefly, norHA HMPs suspended in PBS were centrifuged at 21,000 rcf for 5 minutes. The excess PBS was discarded and 1mM AF tetrazine was added at equal volumes to the HMPs (final concentration 0.5mM AF tetrazine). The mixture was then vortexed to resuspend the HMPs. The reaction was allowed to proceed for 20 minutes, after which the HMP suspension were centrifuged again at 21,000 rcf for 5 minutes. The excess solution was discarded and replaced with fresh PBS. This process was repeated until the solution became clear after centrifuging the HMPs. All scaffolds were fabricated without fiber as described above. The scaffolds were imaged in PBS using a Leica confocal microscope (Stellaris 5). A 20x objective was used and z-stacks of 100 μm was taken for each scaffold.

Detailed explanation of LOVAMAP software has been previously described^37^. Briefly, LOVAMAP uses the location of particles to identify medial axis landmarks within the void space, which are then used as a basis for 3D pore segmentation. 3D pores represent the natural open pockets of space throughout the scaffold that are delineated from one another by narrower regions. “Exterior” pores extend to the surface of the sample, while “interior” pores are completely enclosed by particles and other surrounding pores. Microscope images that were input into LOVAMAP were first processed using previously-described^38^ that reconstructs a 3D scaffold from z-slices, then applies watershed techniques to accomplish particle segmentation. To classify small versus large pores, we again used the largest pore found in 100% norHA scaffolds as the threshold cut-off for small pores.

### Electrospinning norHA fibers

electrospinning solution containing 3.5% w/v norHA, 2.5% w/v 900 kDa polyethylene oxide (PEO, Sigma), 0.05% I2959 (Sigma), and 6.5 mM DTT in deionized water was dissolved overnight. The solution was extruded using a syringe pump at a flow rate of 0.4 mL/hr through a 16G needle. The fibers were collected on an aluminum foil substrate attached to a mandrel spinning at ~1,000 rpm. 13-16 kV positive voltage was applied to the needle and 4kV negative voltage was applied to the collection substrate.

After collection, fibers were crosslinked under UV light for 15 minutes at 10mW/cm^2^ under nitrogen. Once crosslinked, the fibers were wetted with PBS to detach from the collection substrate, then suspended in PBS. The suspension was homogenized at 9,000 rpm for 2 minutes and then filtered through a 40 μm cell strainer (Fisher) to eliminate large aggregations of fibers.

### norHA fiber characterization

fluorescent fibers were made by incorporating 4mg/mL FITC-dextran (1 MDa, Sigma) in the initial electrospinning solution. All other steps were the same to produce fluorescent fibers. After filtering, fibers were diluted 1:1000 in PBS and placed between two glass cover slips. Images were taken using a Leica DMi8 widefield microscope and further dilution of the fibers was carried out until fibers no longer overlap each other in the micrograph. The images were then analyzed in FIJI to find the length and width of each fiber.

### Characterizing degradation of norHA scaffolds

non-fiber scaffolds were fabricated as described above. For fiber-reinforced scaffolds, fibers were first centrifuged at 3,000 rcf to form a pellet. The pellet volume was estimated, and the pellet was resuspended 1:10 fiber to PBS volume ratio. To fabricate the scaffolds, desired volume of fiber was calculated (1%, 5%, or 10%), and corresponding volume of fiber was added to the norHA-gelatin HMP mixture (amount measured as described before for 100%, 75%, and 50%). 10mM LAP and 25mg/mL DTT was then added to the mixture to reach a final concentration of 1mM for each reagent. The rest of the fabrication was carried out in the same way as described for fabricating non-fiber scaffolds.

All scaffolds for this study were fabricated on a glass coverslip substrate. Each coverslip was weighed by itself, and then weighed again with the scaffold on top to find the weight of the scaffold. Each scaffold was then submerged in PBS and cultured in a humidified incubator at 37°C. For each degradation time point, the PBS was removed by pipetting and gently wicking off the remaining PBS with a KimWipe. The weight of coverslip and scaffold was then measured again. Plotting and statistical analysis were done in Rstudio. Statistical comparison was done using ANOVA and Tukey’s post-hoc test.

### Rheological tests for HMP scaffolds

Time sweeps were done for HMP scaffolds to assess their mechanical properties. Scaffolds were fabricated into disks with 1mm height and 8mm diameter. This is done by first biopsy punching a 1mm thick PTFE sheet (McMasters) to create an 8mm diameter opening. The HMPs were loaded into the opening, then a glass slide is placed on top to flatten the HMPs so that they conform to the opening. Excess HMPs that overflowed between the glass slide and the PTFE sheet were simply wiped away after the glass slide was removed. The HMPs were then photo-crosslinked for 1 minute under 10mW/cm^2^ UV light.

Rheology was conducted on a Peltier plate temperature controlled at 20°C. The fabricated 8mm scaffolds were loaded onto the Peltier plate and positioned under an 8mm sandblasted parallel plate geometry. Each scaffold is conditioned by applying a small axial force (0.3-0.8 N) prior to oscillatory testing. This accounts for the potential uneven topography produced during scaffold fabrication and ensures that all scaffold is sufficiently contacting the geometry. Time sweeps were done at 1% strain and 1 Hz frequency for 60 seconds.

To assess mechanical properties of scaffolds post gelatin HMP sacrifice, after time sweep was done for the pre-sacrifice scaffolds, the scaffolds were submerged in PBS and cultured for 20 minutes at 37°C. After culturing, the scaffolds were gently washed with excess PBS, also at 37°C, to remove liquefied gelatin. The same time sweep with conditioning was conducted again on the scaffolds. Statistical analysis was done using paired t-tests for these samples.

### HUVEC culture in fiber-reinforced, porous GHs

HUVECs (Lonza) was cultured in EGM-2 medium (Lonza) at 5% CO2 and 37°C in a humidified environment. HUVECs Passage 4-8 was used for this experiment. HUVECs were detached from plate using 0.05% trypsin, centrifuged at 300 rcf for 3 minutes, and resuspended in EGM-2 at ~50 million cells per mL.

RGD-modified norHA HMPs were made by mixing the norHA HMPs with RGD and LAP to reach a final concentration of 1mM RGD and 1mM LAP (norHA HMPs comprise half the volume of the solution). This mixture was placed under UV light at 10mW/cm^2^ for 5 minutes, then washed twice with PBS to eliminate excess RGD and LAP.

The RGD-norHA HMPs were mixed at desired ratios with sacrificial gelatin HMPs along with LAP (1mM final concentration) and DTT (2mM final concentration). The mixture was jammed by centrifugation at 21,000 rcf for 5 minutes. The dense HUVEC suspension was then mixed with the HMPs to reach a final density of 5 million cells/mL. The mixture was transferred to a confocal plate and subsequently UV crosslinked at 15 mW/cm^2^ for 1 minute and cultured for 4 days in EGM-2 media (Lonza).

To stain the HUVEC-containing scaffolds, the scaffolds were fixed by submerging in 4% paraformaldehyde. After fixation, permeation was carried out using 0.1% Triton X-100. The scaffolds were then blocked with 3% BSA in PBS and stained with fluorescent Phalloidin (Thermo Fisher) overnight. After PBS wash, DAPI (Thermo Fisher) staining was carried out for 30 minutes and the scaffolds were washed again and imaged using a confocal microscope (Stellaris 5).

### Injection of HUVEC and HMPs

HUVECs (Lonza) was cultured in EGM-2 medium (Lonza) at 5% CO2 and 37°C in a humidified environment. HUVECs Passage 6-8 was used for this experiment. HUVECs were detached from plate using 0.05% trypsin, centrifuged at 300 rcf for 3 minutes, and resuspended in EGM-2 at 30 million cells per mL.

RGD-modified norHA HMPs were made by mixing the norHA HMPs with RGD and LAP to reach a final concentration of 1mM RGD and 1mM LAP (norHA HMPs comprise half the volume of the solution). This mixture was placed under UV light at 10mW/cm^2^ for 5 minutes, then washed twice with PBS to eliminate excess RGD and LAP.

The RGD-norHA HMPs were centrifuged at 21,000 rcf for 5 minutes. The dense HUVEC suspension was then mixed with the HMPs to reach a final density of 2 million cells/mL. The mixture was then loaded into a 1 mL syringe and subsequently extruded onto a 6-well cell culture plate through a bevel, 18G needle. The material was subsequently UV crosslinked at 15 mW/cm^2^ for 1 minute.

For live/dead staining, HUVECs-HMPs mixture were collected prior to loading the material into the syringe, while post-printing mixtures were collected either immediately after extrusion or after 3 days of culture. The cells were stained using a Live/Dead viability kit (L3224, Invitrogen) and imaged in a plastic-bottom culture plate by a Leica DMi8 widefield microscope. Statistical analysis was performed using GraphPad Prism 9. Statistical comparison was made using one-way analysis or variance (ANOVA) with Tukey post hoc test.

## Notes

### Competing Interest Statement

The authors have declared no competing interest.

### Summary of Updates

Changed corresponding author designation. It should now be Dr. Chris Highley as the corresponding author, from Dr. Jack Whitewolf being the corresponding author in the previous submission.

