## Supplemental Data for "Highly porous granular hydrogels reinforced by electrospun hydrogel fibers for long-term stability"

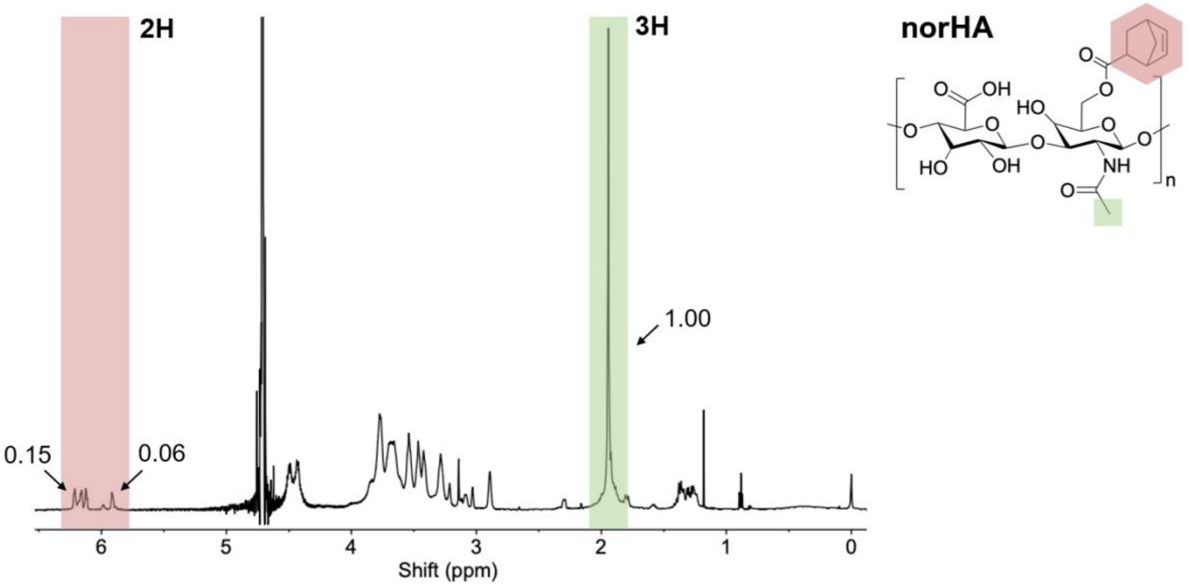


**Figure S1**. [1]H NMR spectrum of norbornene-modified HA with 0.3 degree of substitution.

### Table S1. Varying norHA HMP compositions and available reactive groups

| | Soft | Medium | Stiff |
|---|---|---|---|
| norHA (wt%) | 2 | 3 | 4 |
| DTT (mM) | 2 | 3 | 4 |
| Total norbornene (mM) | 15 | 22.5 | 30 |
| Remaining norbornene (mM) | 11 | 16.5 | 22 |
| % norbornene consumed | 26.7 | 26.7 | 26.7 |
| | | norHA DoS: | 0.3 |

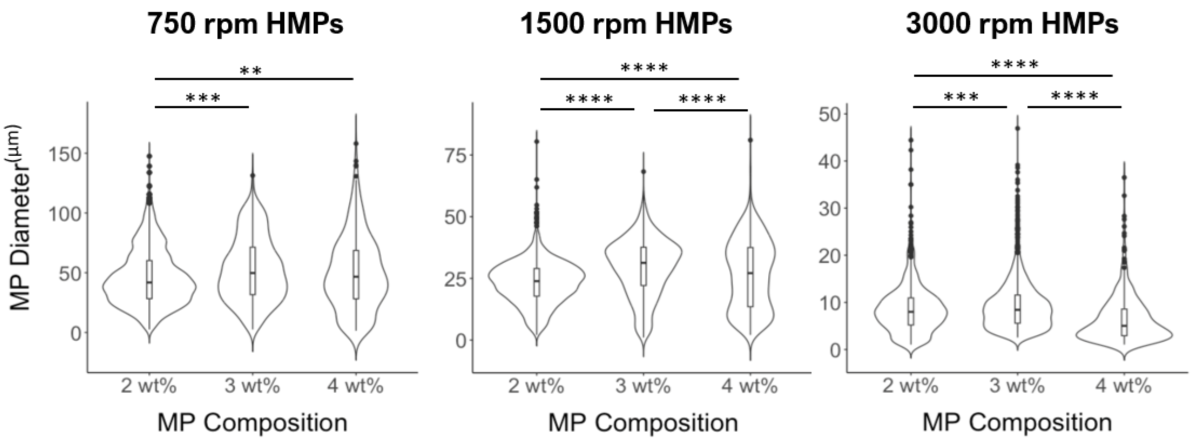


**Figure S2**. Size distributions comparing norHA HMPs with varying compositions. Effect sizes were also calculated for each comparison. The largest effect size calculated was 0.55, accounting for a medium effect (>0.5), all other comparisons had a small effect (>0.2) or less.

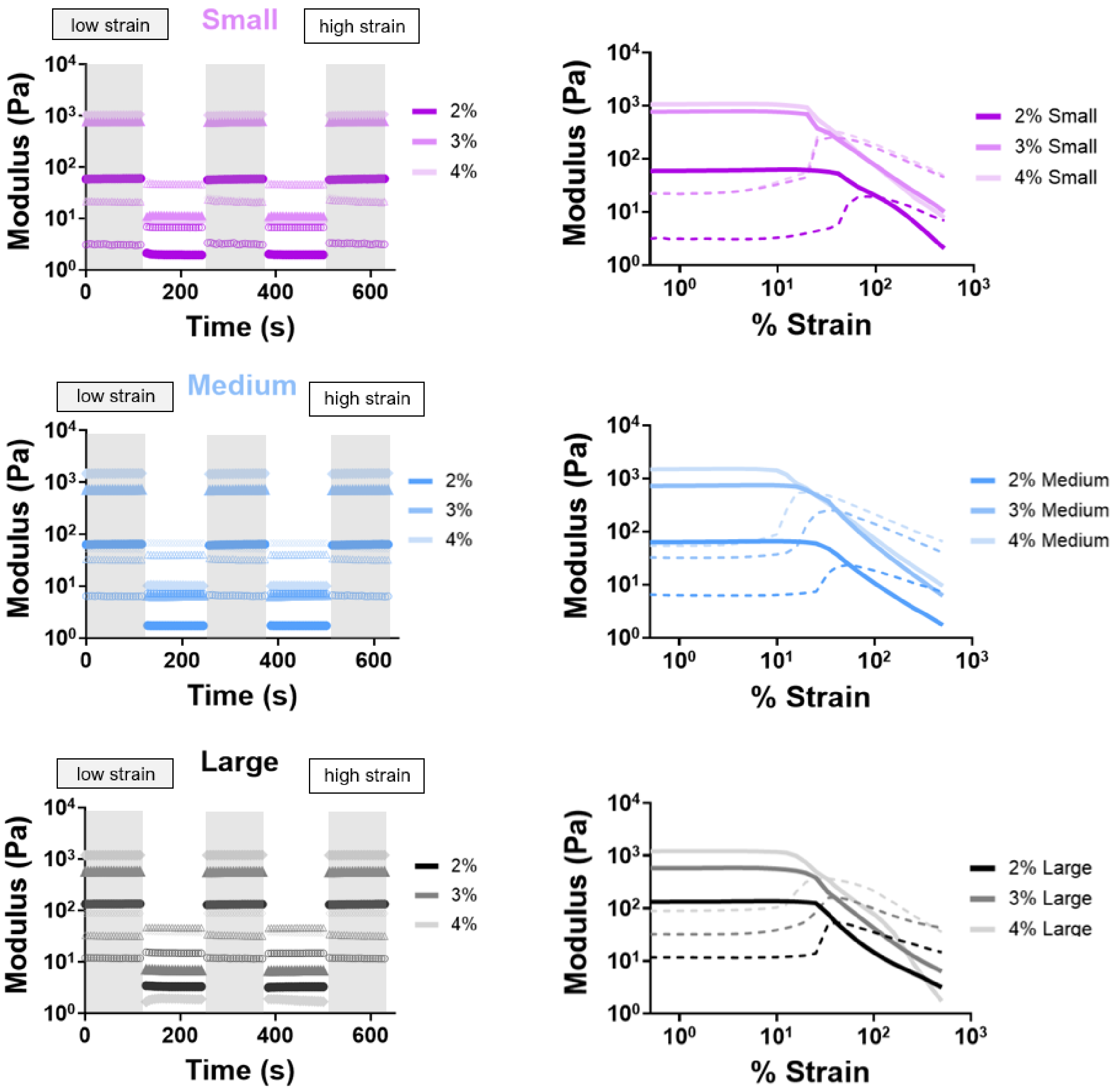


**Figure S3**. Representative rheology results for all norHA HMP compositions and sizes. *Left column:* time sweeps with strains alternating from high (500%) strain to low (1%) strain. *Right column:* strain sweeps from 0.1% strain to 500% strain. From top to bottom: rheology results of small, medium, and large norHA HMPs, respectively.

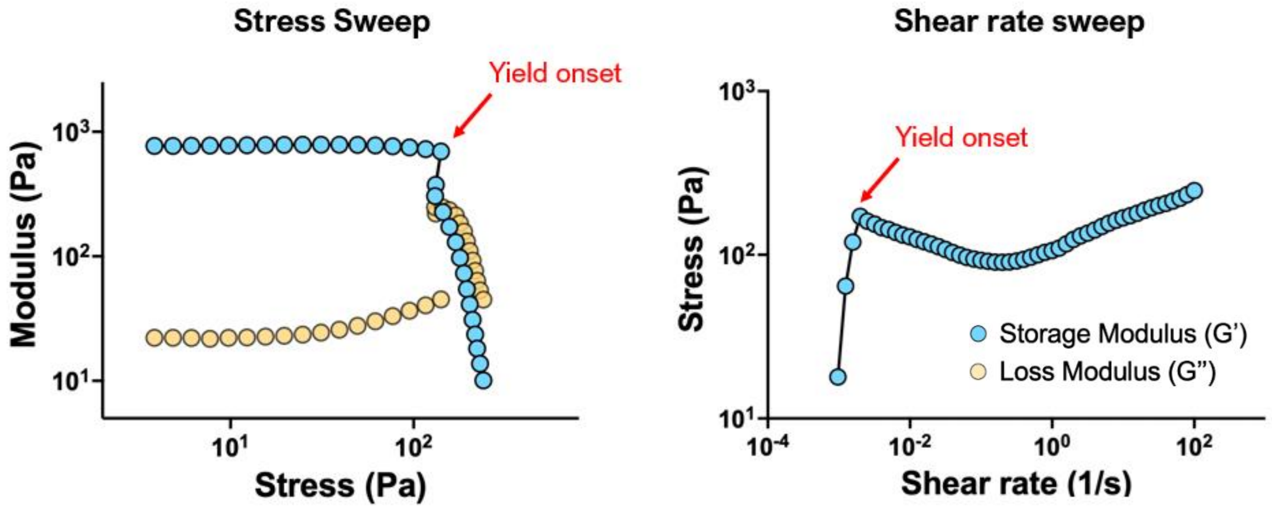


**Figure S4**. Representative rheology curves for packed norHA HMP stress sweep (left) and shear rate sweep (right). Red arrow points to representative data points used for quantifying and confirming yield stress.

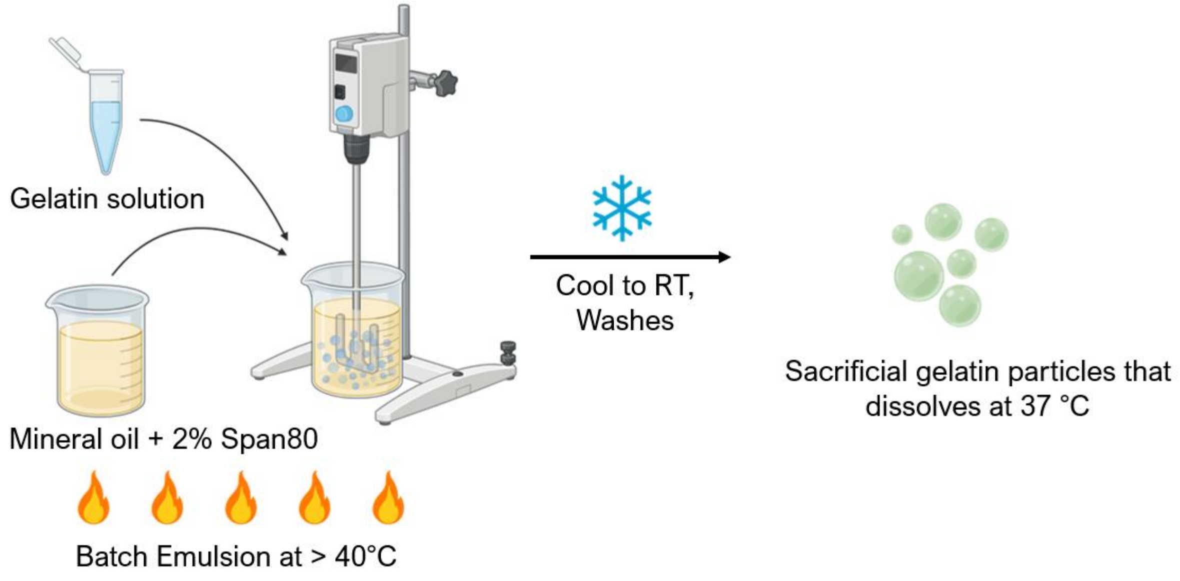


**Figure S5**. Gelatin HMP fabrication scheme. Gelatin solution was homogenized at varying speeds in oil and surfactant at 40 degrees Celsius. The mixture is then cooled to room temperature and washed in the same method as norHA HMPs to obtain gelatin HMPs.

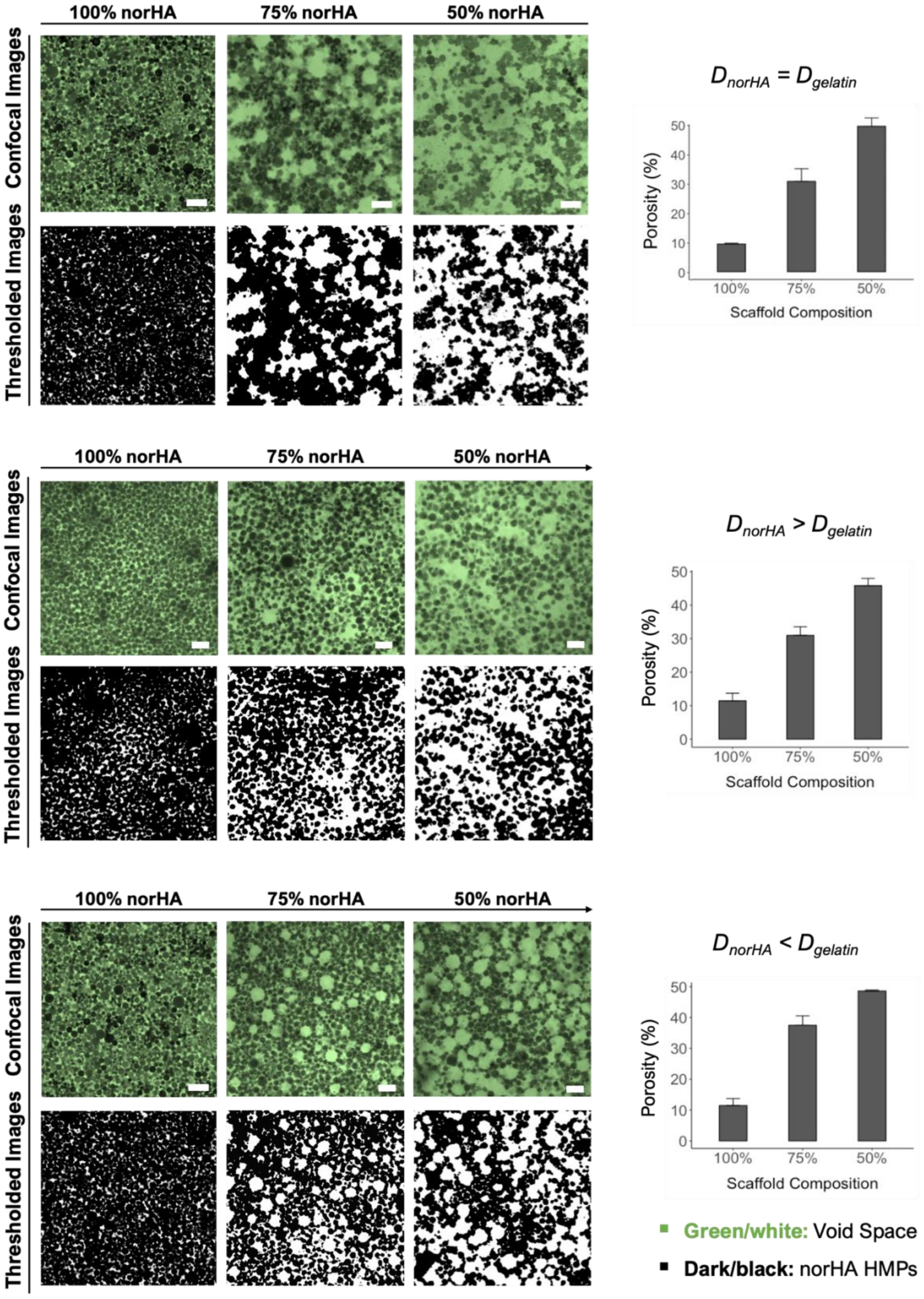

100% norHA
75% norHA
50% norHA
Confocal Images
Thresholded Images
$D_{norHA} = D_{gelatin}$
Porosity (%)
Scaffold Composition
100%
75%
50%
$D_{norHA} > D_{gelatin}$
$D_{norHA} < D_{gelatin}$
Green/white: Void Space
Dark/black: norHA HMPs

**Figure S6.** From to top to bottom: 100% norHA, 75% norHA, and 50% norHA scaffolds created with gelatin HMPs with diameters greater than (top), equal to (middle), or less than (bottom) the diameters of norHA HMPs. Particles are black in both confocal images and thresholded images. Void space is shown in green in confocal images and white in thresholded images. Quantification of void space is shown to the right. Scale bars = 50 µm. Error bars denote standard deviations.

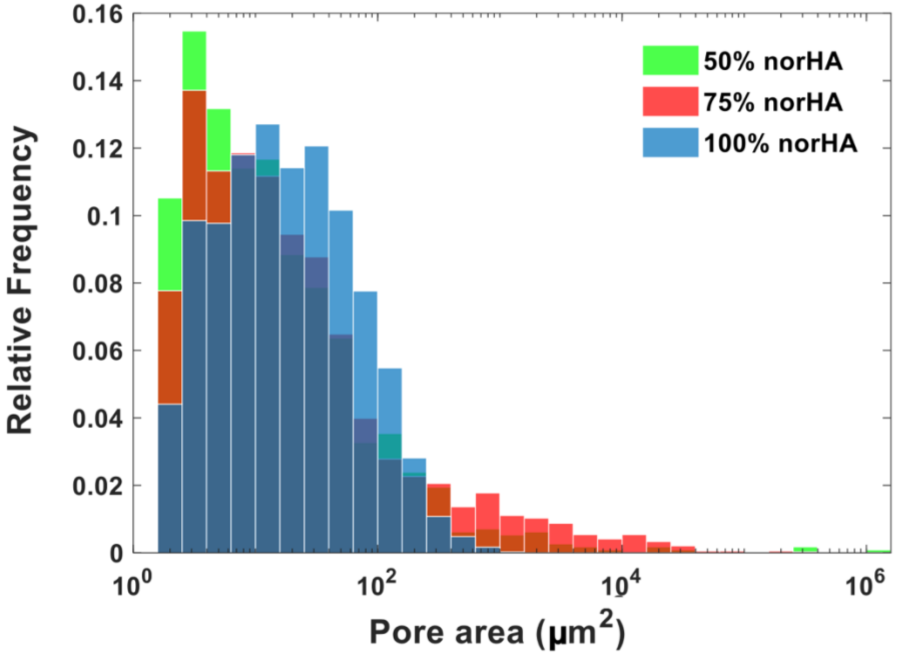


**Figure S7.** Number-based pore area distribution for each scaffold composition. For each scaffold group (50%, 75%, and 100% norHA), pores from n = 3 scaffolds were sampled to generate the distribution.

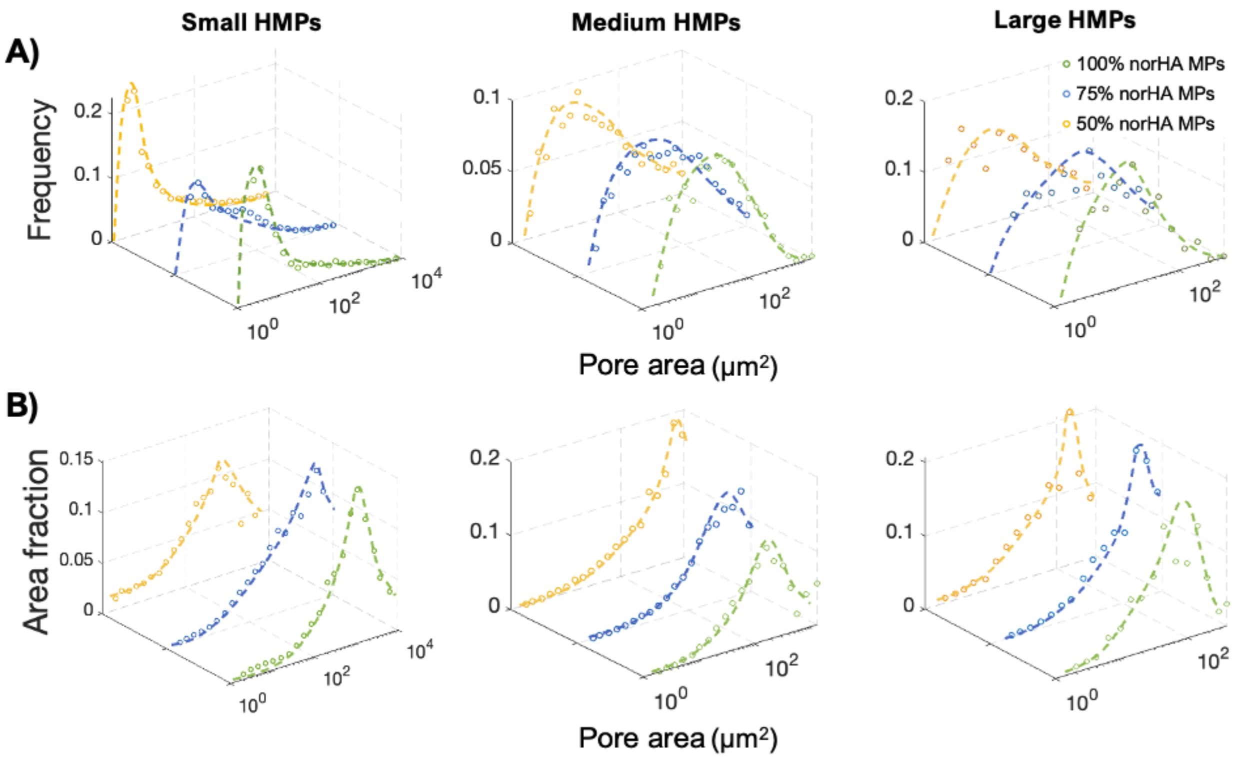


**Figure S8.** Pore area distributions using only small pores. For each HMP size, Small pores are determined using the largest pore found in 100% norHA scaffolds as the cut-off point. **A)** Pore area distribution based on the number of pores. **B)** Pore area distributions based on the area fraction of pores

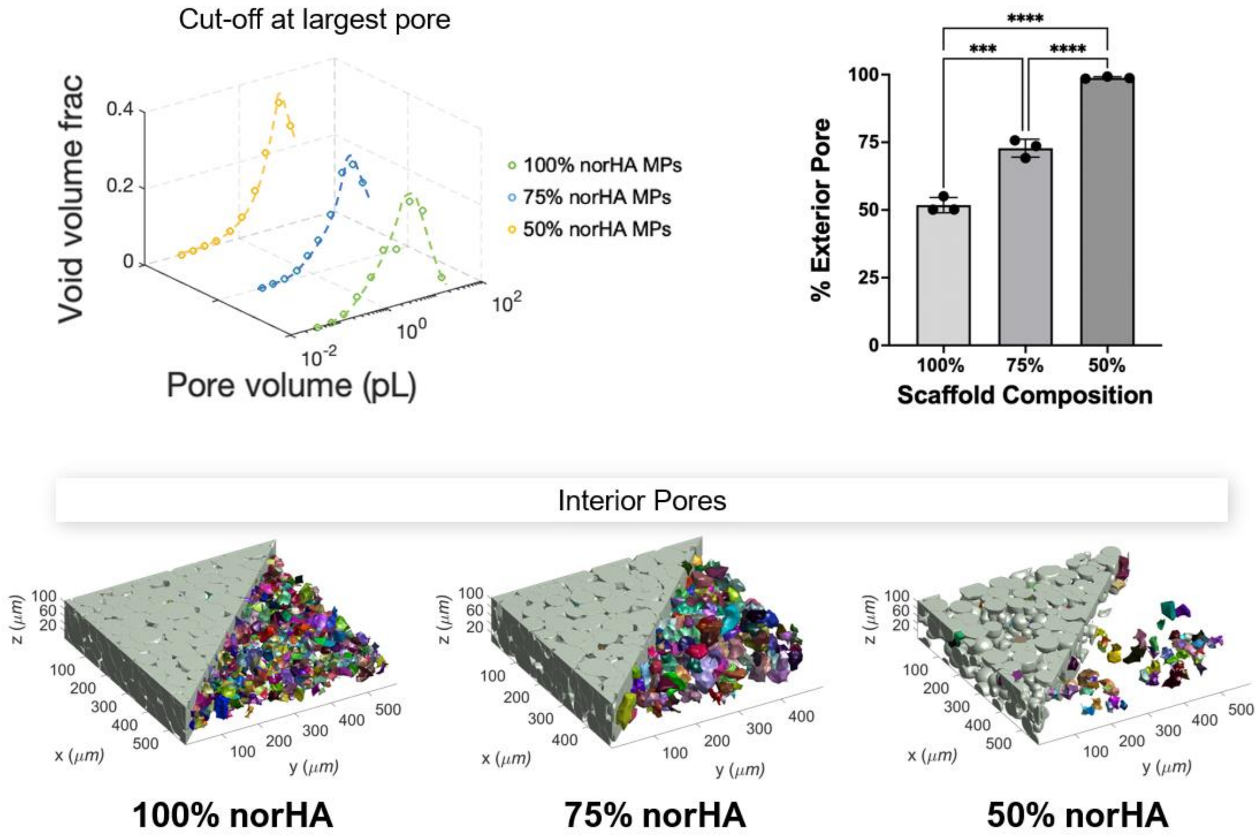


**Figure S9.** *Top left:* volume fraction-based pore volume distributions using the largest pore in 100% norHA scaffolds as the cut-off point. *Top right:* percent of exterior pores for each scaffold composition. *Bottom:* representative LOVAMAP-generated scaffolds with only interior pores shown. Error bars denote standard deviations.

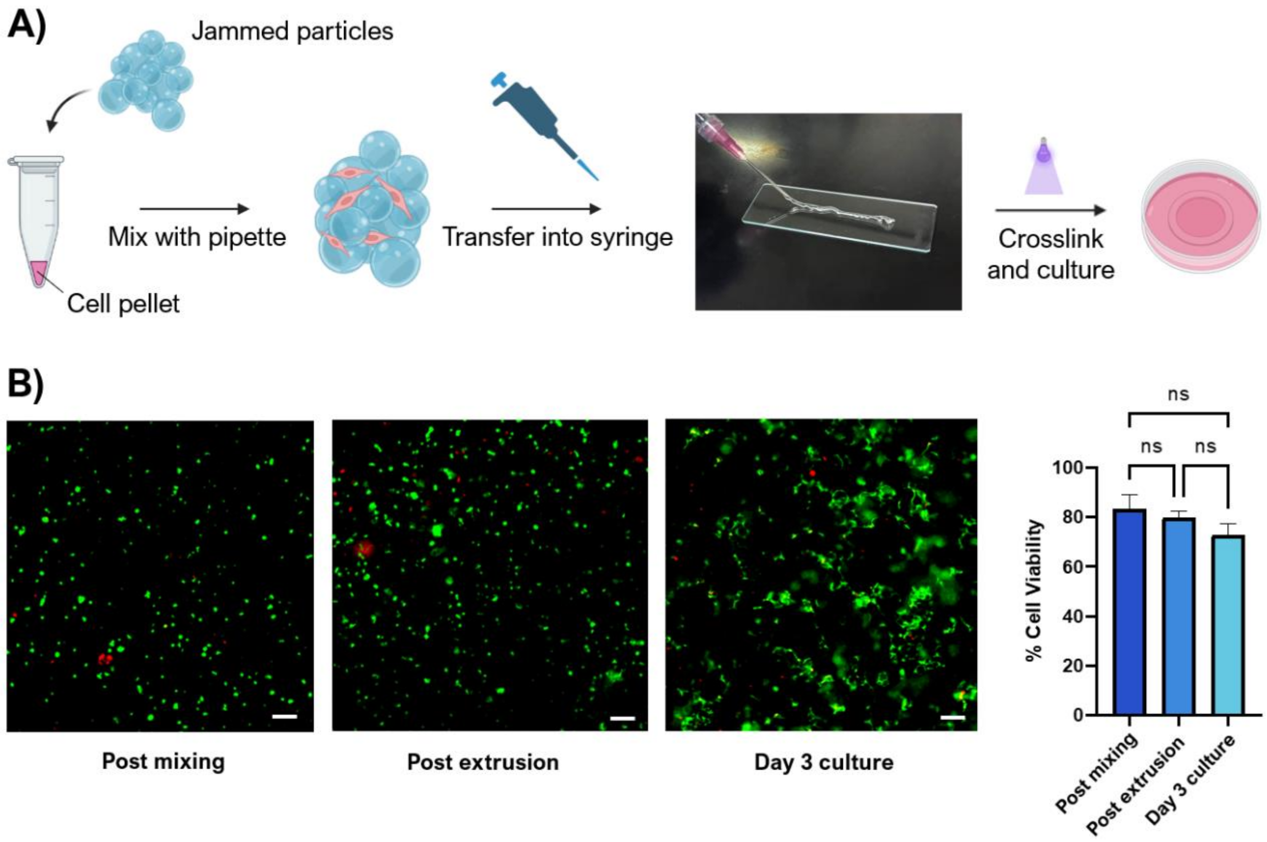


**Figure S10. A)** The process for HUVEC incorporation, extrusion, and scaffold formation. **B)** Representative images of live/dead stain carried out independently after HUVECs are mixed with HMPs (left), after the cells-HMPs mixture are extruded through the syringe (mid), and after 3 days of culture (right). **C)** Quantified cell viability for each timepoint in panel B. Error bars denote standard deviations.
